# Efficient near telomere-to-telomere assembly of Nanopore Simplex reads

**DOI:** 10.1101/2025.04.14.648685

**Authors:** Haoyu Cheng, Han Qu, Sean McKenzie, Katherine R. Lawrence, Rhydian Windsor, Mike Vella, Peter J. Park, Heng Li

## Abstract

Telomere-to-telomere (T2T) assembly is the ultimate goal for de novo genome assembly. Existing algorithms capable of near T2T assembly all require Oxford Nanopore Technologies (ONT) ultra-long reads which are costly and experimentally challenging to obtain and are thus often unavailable for samples without established cell lines. Here, we introduce hifiasm (ONT), the first algorithm that can produce near T2T assemblies from standard ONT Simplex reads, eliminating the need for ultra-long sequencing. Compared to existing methods, hifiasm (ONT) reduces the computational demands by an order of magnitude and reconstructs more chromosomes from telomere to telomere on the same datasets. This advancement substantially broadens the feasibility of T2T assembly for applications previously limited by the high cost and experimental requirement of ultra-long reads.

---

With the advent of accurate long reads, notably Pacific Biosciences High-Fidelity (PacBio HiFi) reads^1^, the new generation of assembly algorithms has revolutionized de novo assembly^2–4^. For diploid genomes, these tools routinely produce haplotype-resolved assemblies, accurately reconstructing both haplotypes^5^, but they often struggle with centromeres or long segmental duplications due to the limited read lengths of PacBio HiFi at 10–20 kilobases (kb). To achieve near T2T human assemblies where each chromosome is completely resolved from end to end^6^, existing assemblers including Verkko^7, 8^and hifiasm (UL)^9^ have to rely on ONT ultra-long reads^10, 11^ of *≥* 100 kb to assemble through regions failing HiFi reads^12^. However, generating ultra-long reads is costly and demands large amount of high molecular weight DNA at tens of micrograms per human sample^13^, ~40 times higher than the DNA input with the standard ONT protocol. As a result, ultra-long reads are rarely produced for clinical specimens or by biodiversity projects. This greatly limits the practicality of near T2T assembly.

There is an urgent need to develop new algorithms to improve the accessibility of T2T genome assembly. The longer read lengths and rapidly improving accuracy of ONT reads offer the potential for achieving T2T assembly using ONT as the sole long-read sequencing technology. Current ONT sequencing protocols produce two types of reads: Duplex and Simplex. Duplex reads are as accurate as PacBio HiFi reads and work as well in assembly^14^but they are expensive to generate and rarely available. Therefore, ONT sequencing primarily focuses on Simplex reads^13^, which are longer in length and cheaper to produce. Nevertheless, the de novo assembly of Simplex reads remains challenging due to their higher non-random, recurrent sequencing error rates^15^. These errors conflict with the key assumption used in haplotype-resolved assembly algorithms, such as hifiasm^2, 5, 9^, HiCanu^3^, LJA^4^, and Verkko^7, 8^, that sequencing errors are random. Currently, ultra-long ONT sequencing can generate only Simplex reads. Therefore, in this study, the term “ultra-long” specifically refers to ultra-long Simplex reads.

Several assembly workflows have been proposed to generate haplotype-resolved assemblies using ONT Simplex reads ^16–19^. They first build a consensus assembly by collapsing multiple haplotypes and then reconstruct each haplotype through the decompression from the consensus. This approach may fail in highly heterozygous regions and complex repetitive regions that cannot be accurately represented in the initial consensus assembly^20^. A recent error-correction tool called HERRO^21^ leverages deep learning to correct ONT Simplex reads before feeding them into existing assemblers like hifiasm and Verkko. While promising, HERRO is computationally intensive and demands high-end GPUs. Furthermore, while HERRO has been extensively validated using ultra-long Simplex reads, it has not been shown to achieve near T2T assembly with standard ONT Simplex reads alone. HERRO is not an answer to near T2T assembly at the population scale.

To address the practical limitations of existing assembly methods, we developed hifiasm (ONT) to assemble the widely 1 used ONT R10.4.1 standard Simplex reads without ultra-long sequencing. It introduces a fast error correction algorithm that leverages read phasing to overcome the higher recurrent error rate of ONT Simplex reads. On real data, hifiasm (ONT) often assembles multiple chromosomes from telomere to telomere with substantial reduction in time, labor, and cost in comparison to the current state of art.

## Results

### Hifiasm (ONT) algorithm for ONT Simplex reads

Current assemblers optimized for PacBio HiFi reads all have an error correction step to correct HiFi reads to nearly error-free. This step assumes the errors are rare and random (Fig. 1a), which is approximately true for PacBio reads. However, the assumption does not hold for ONT Simplex reads. In comparison to HiFi reads, ONT reads have higher error rates and the ONT sequencing errors tend to be recurrent – the same error may occur in multiple reads at the same genomic position (Fig. 1b). This makes it challenging to distinguish ONT sequencing errors from true heterozygous variants. As a result, error correction algorithms in current HiFi assemblers do not work well with ONT Simplex reads.

**Figure 1.**
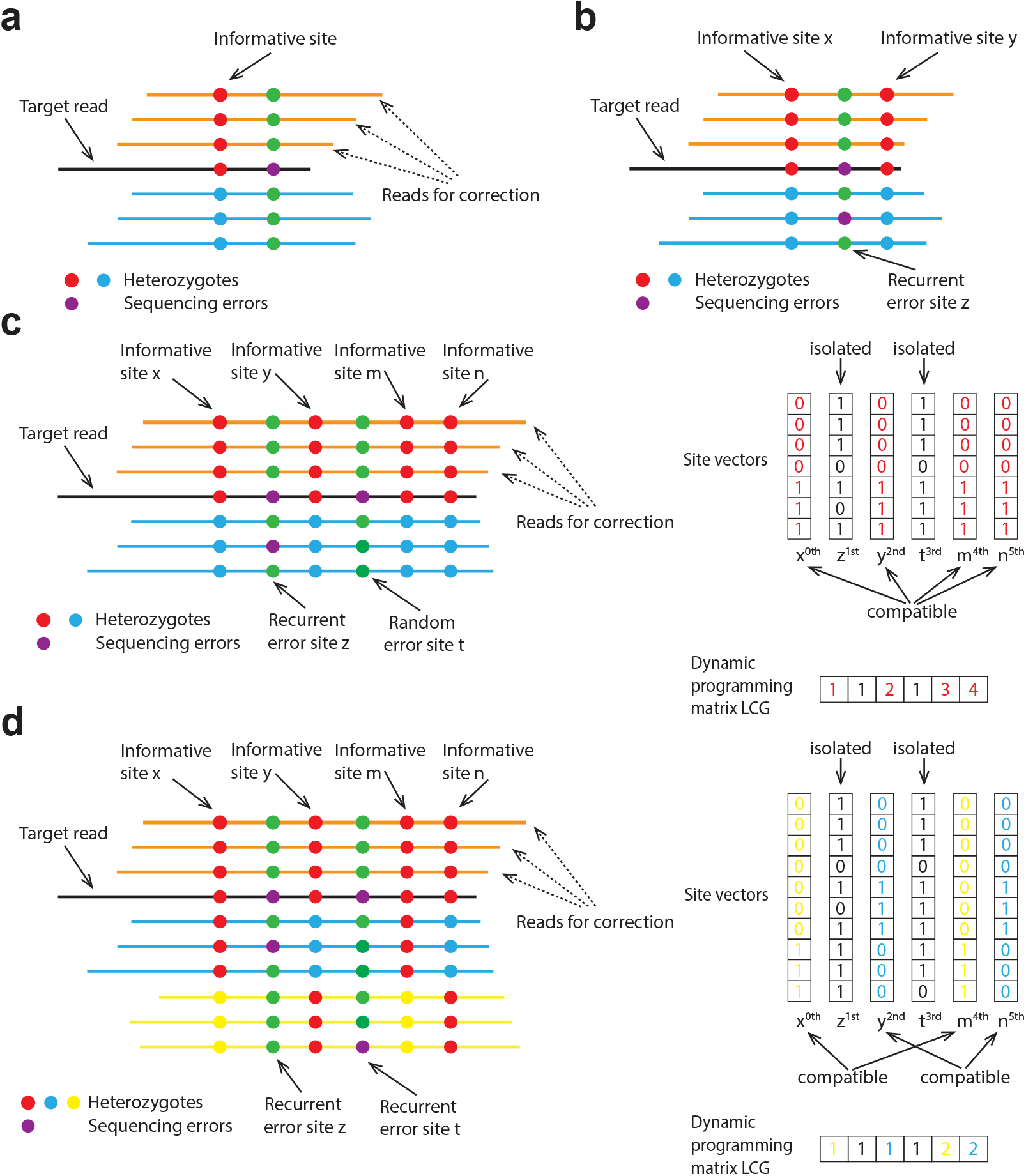
Error correction of ONT Simplex reads. **(a)** Error correction in existing hifiasm for PacBio HiFi reads. Hifiasm identifies informative sites where each allele (represented in red or blue) is supported by multiple reads. Sequencing errors are represented by purple dots. The algorithm then corrects the target read (black) using supporting reads (orange) that match the target read across all informative sites. **(b)** Recurrent sequencing errors in ONT Simplex reads. The existing error-correction approach in hifiasm incorrectly identifies recurrent sequencing errors as informative sites (illustrated as purple dots) as it is supported by two reads. In this case, all blue reads are correctly excluded due to real informative sites (*x* and *y*), while all orange reads are mistakenly discarded because of the false-positive informative site *z*. With existing error correction approach, no reads remain available for correction. **(c)** Error correction in hifiasm (ONT) with two haplotypes. The site vectors corresponding to informative sites *x, y, m*, and *n* (highlighted in red) are identified as mutually compatible. These sites can be grouped into the same cluster using the dynamic programming matrix. Sites resulting from sequencing errors, such as *z* and *t*, are incompatible with other sites and remain unclustered. **(d)** Error correction in hifiasm (ONT) with more than two haplotypes or repeat copies. The target read (black) and the orange reads originate from haplotype/repeat copy 1, the blue reads from haplotype/repeat copy 2, and the yellow reads from haplotype/repeat copy 3. Using the dynamic programming matrix, sites *x* and *m* can be grouped into one cluster (highlighted in yellow), while sites *y* and *n* form another cluster (highlighted in blue). Sites *x* and *m* exclude reads from haplotype/repeat copy 3, whereas sites *y* and *n* exclude reads from haplotype/repeat copy 2.

Hifiasm (ONT) overcomes the limitation of current methods by exploiting the phasing of long reads: a true heterozygous site is in phase with nearby heterozygotes sites but a site loaded with recurrent sequencing errors is not (Fig. 1c, d). It employs a dynamic programming based algorithm for joint phasing and the identification of sequencing errors and it considers base quality scores as well. With the new algorithm, hifiasm (ONT) can correct most ONT Simplex reads to error-free. It also introduces other improvements to the assembly step in comparison to earlier versions (Online Methods).

### Seven human genomes sequenced using ONT standard Simplex reads

To demonstrate the capabilities of hifiasm (ONT), we generated standard ONT Simplex reads for seven human samples and thoroughly evaluated hifiasm (ONT)’s performance. These samples, including HG001, HG002, HG003, HG004, HG005, HG006, and HG007, are well-characterized by the Genome in a Bottle (GIAB) Consortium for benchmarking^25^. Each sample was sequenced using ONT Simplex sequencing with one or two R10.4 flow cells, targeting to produce non-ultra-long reads with approximately 50× or higher coverage per genome (Online Methods). The average read N50 of these datasets is 30 kb, defined as the length of the shortest read that cumulatively covers 50% of the total read size (Supplementary Fig. 1). This value is often used as an indicator of the average read length.

As shown in Supplementary Fig. 1b, ONT standard Simplex reads are generally longer than the PacBio HiFi reads we collected for the same samples. Moreover, ONT Simplex sequencing yields a broader read length distribution than PacBio HiFi, increasing the likelihood of obtaining reads that are substantially longer than the average (Supplementary Fig. 1a). Such long reads are particularly valuable for resolving complex and repetitive regions during assembly^26^. Compared to ONT ultra-long reads (typically with an N50 *>* 100 kb), ONT standard Simplex reads are shorter; however, they offer several practical advantages, including higher throughput, lower cost, and up to 40-fold reduction in DNA input requirements^11, 13^. These benefits make ONT standard sequencing more accessible, especially for sample types where ultra-long protocols are not feasible.

### Benchmarking human genome assembly with ONT standard Simplex reads

We compared hifiasm (ONT) algorithm from the hifiasm toolkit to another contemporary T2T assembler Verkko^7, 8^. For HG001, HG002, and HG005 with parental data for phasing, we performed trio-binning assembly^27^using both hifiasm (ONT) and Verkko for comparison. Since Verkko cannot directly assemble ONT Simplex reads due to their higher error rates, we first preprocessed the reads with HERRO^21^for error correction before assembling them with Verkko. To evaluate the remaining samples without parental data, we also evaluated assembly quality using hifiasm (ONT) in its dual-assembly mode^5^, which can generate high-quality assemblies using ONT reads alone. Unlike trio-binning assemblies that are fully phased, ONT-only dual assemblies are partially phased, relying on homologous information between the two haplotypes rather than additional data such as Hi-C or parental information. Despite being only partially phased, a dual assembly still represents a complete diploid genome and is highly contiguous. Verkko was not used for samples without parental data since it does not support dual-assembly mode. All hifiasm assemblies were performed on a standard computational server with 64 CPUs, whereas Verkko+HERRO was run on a combination of different servers optimized for performance, utilizing more than 64 CPUs and multiple GPUs (see Supplementary Section 1.4 for more details).

As shown in Table 1, hifiasm is approximately an order of magnitude faster than Verkko+HERRO when assembling ONT Simplex reads generated using the Super Accurate (SUP) basecalling model. In addition, unlike HERRO requiring high-end GPUs for error correction, hifiasm is an all-in-one toolkit capable of assembling directly from raw reads using only CPUs. This further reduces computational costs and simplifies the deployment. Moreover, hifiasm assemblies consistently exhibit higher quality than those produced by Verkko+HERRO. Using ONT standard Simplex reads, hifiasm successfully reconstructs 9–22 chromosomes from telomere to telomere across different samples, while Verkko+HERRO fails to produce any complete T2T contigs, except for HG001 with two T2T contig. This advantage of hifiasm is also evident in assembly contiguity, particularly in the contig N50. We noted that Verkko+HERRO slightly outperforms hifiasm in terms of QV scores, which reflect per-base assembly accuracy (Table 1 and Supplementary Table 1). To investigate this, we compared the HG002 assemblies generated by both tools against the HG002 Q100 reference^28^. We found that most per-base errors unique to the hifiasm assembly originated from long homopolymer regions (see Supplementary Fig. 2 and Supplementary Table 6). High error rates in long homopolymer regions are a known limitation of ONT reads^29^, making these errors challenging to correct through assembly algorithms. We observed large numbers of sequencing errors remaining within long homopolymer regions in both hifiasm and Verkko+HERRO assemblies, although Verkko+HERRO performed slightly better. Fully resolving these errors would require an additional polishing step, either using PacBio HiFi reads with higher base accuracy, or re-using the ONT Simplex reads with a polisher such as Medaka or Dorado Polish, which is more computationally intensive.

**Table 1.**
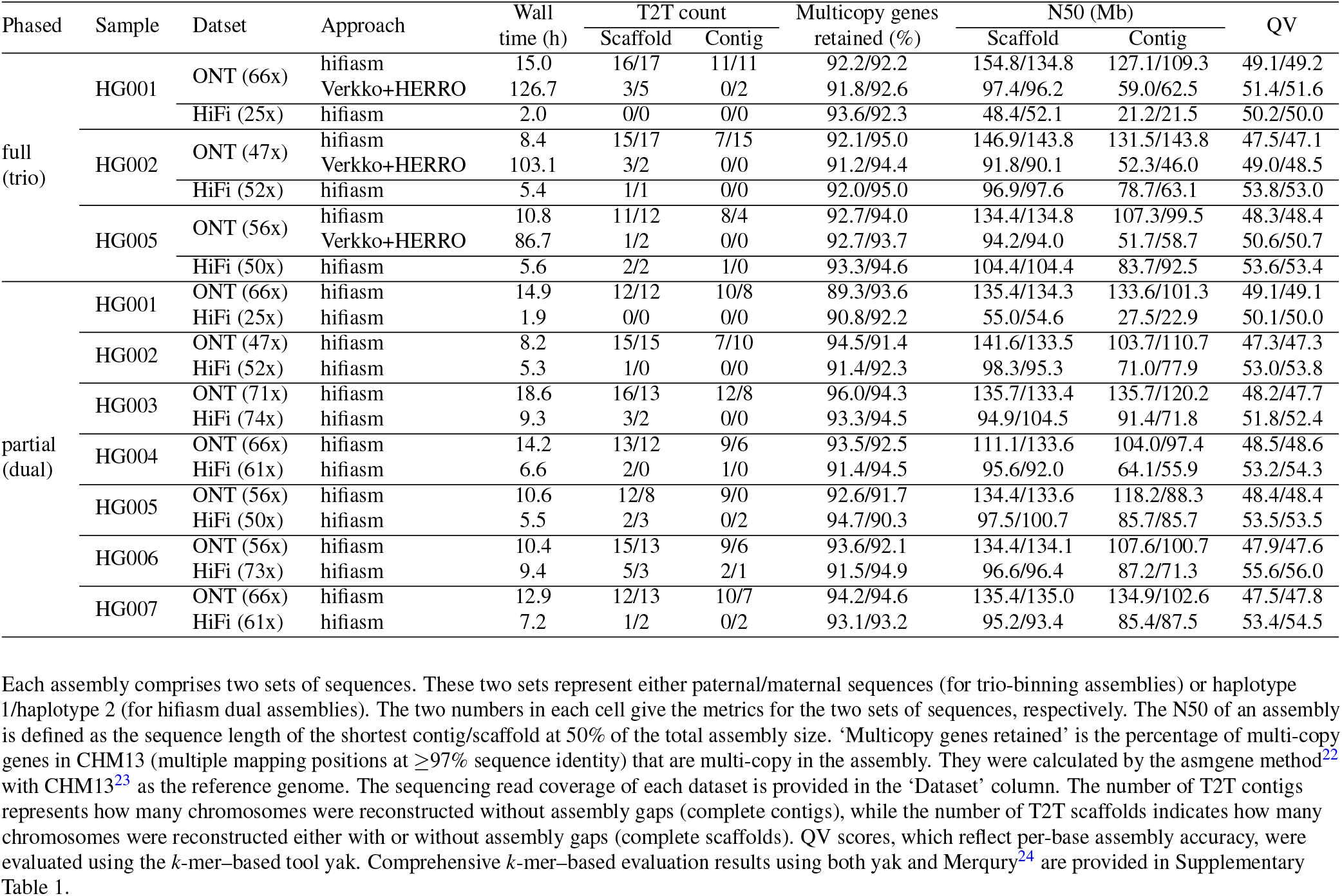
Statistics of different assemblies using ONT standard Simplex (SUP basecalling model) and PacBio HiFi reads.

In addition to the assembly results shown in Table 1, which use ONT reads basecalled with the SUP model, we also tested hifiasm (ONT) assemblies using reads produced by the High Accuracy (HAC) model (Supplementary Table 2). Compared to the SUP model, the HAC model is about 10 times faster in basecalling on an NVIDIA H100 GPU, but at the expense of lower base accuracy^30^. Nonetheless, for each sample, hifiasm (ONT) achieved comparable assembly quality using the less accurate HAC reads, with QV scores approximately 3 points lower on the Phred scale. These QV scores can be further improved by applying a post-assembly polishing tool like Dorado Polish, which is designed to work with both HAC and SUP reads and can substantially improve the base-level accuracy of the draft assembly.

### Comparison of assemblies produced by ONT standard Simplex and PacBio HiFi Reads

We also compared assemblies generated from ONT standard Simplex reads and PacBio HiFi reads. For fair comparison, we produced HiFi-based assemblies for each sample at similar coverage levels, except for HG001 due to insufficient publicly available HiFi data. As illustrated in Table 1, ONT assemblies exhibit substantially higher contiguity, as indicated by their N50 values and the number of T2T contigs/scaffolds. This superior contiguity mainly results from ONT standard reads being tens of kilobases longer than PacBio HiFi reads. Evaluating assembly quality within repetitive regions by examining multicopy gene retained rates, we found that ONT assemblies show quality comparable to PacBio HiFi assemblies across all samples. This demonstrates that despite high sequencing error rates in ONT Simplex reads, our algorithm effectively corrects these errors while avoiding the common problem of collapsing highly similar repeats observed in existing ONT assemblers^20^ like Napu (Shasta)^18^(see Supplementary Section 6). We further compared the number of annotated immunological genes per human genome assembly^32^ (see Supplementary Fig. 3). These genes, including the HLA and KIR loci, are typically repetitive and multi-allelic, further supporting that ONT assemblies achieve a quality comparable to PacBio HiFi assemblies. As anticipated, ONT assemblies had lower QV scores than PacBio HiFi assemblies, primarily due to persistent sequencing errors in long homopolymer regions, which are challenging to fully correct. The lower per-base accuracy of ONT assemblies also results in slightly higher phasing switch and hamming error rates compared to PacBio HiFi assemblies (Supplementary Table 1). In terms of computational performance, ONT assemblies typically require 1.5 to 2 times longer than HiFi assemblies at similar coverage levels; however, most ONT assemblies can be completed within approximately half a day using 64 CPUs.

### Assembling using ONT ultra-long reads

We then evaluated the performance of hifiasm (ONT) and Verkko+HERRO using ONT ultra-long reads (Fig. 2). To ensure a comprehensive comparison, we tested two diploid human genomes (HG002 and HG02818) from the Human Pangenome Reference Consortium (HPRC)^33^and three haploid non-human genomes: *Arabidopsis thaliana* (Arabidopsis)^21^, *Danio rerio* (zebrafish)^34^, and *Solanum lycopersicum* (tomato)^14^. Consistent with previous results, hifiasm (ONT) remained an order of magnitude faster than Verkko+HERRO and did not require GPU resources. Although Verkko+HERRO assemblies improved substantially with ultra-long reads, hifiasm (ONT) still outperformed in all quality metrics except QV (Fig. 2 and Supplementary Table 1). For most samples, hifiasm (ONT) successfully reconstructed the majority of chromosomes from telomere to telomere: 41 out of 46 for HG002, 44 out of 46 for HG02818, 3 out of 5 for Arabidopsis, and 21 out of 25 for zebrafish. Tomato is an exception, likely due to its lower sequencing coverage (33×), compared to 68× for HG002, 58× for HG02818, 153× for Arabidopsis, and 145× for zebrafish. To further assess assembly quality in non-human genomes, we additionally tested the ONT assembly of *Linum usitatissimum* (flax)^35^(see Supplementary Table 4). Despite relying only on ONT standard reads, hifiasm (ONT) still outperformed Verkko+HERRO, reconstructing many chromosomes completely from telomere to telomere.

**Figure 2.**
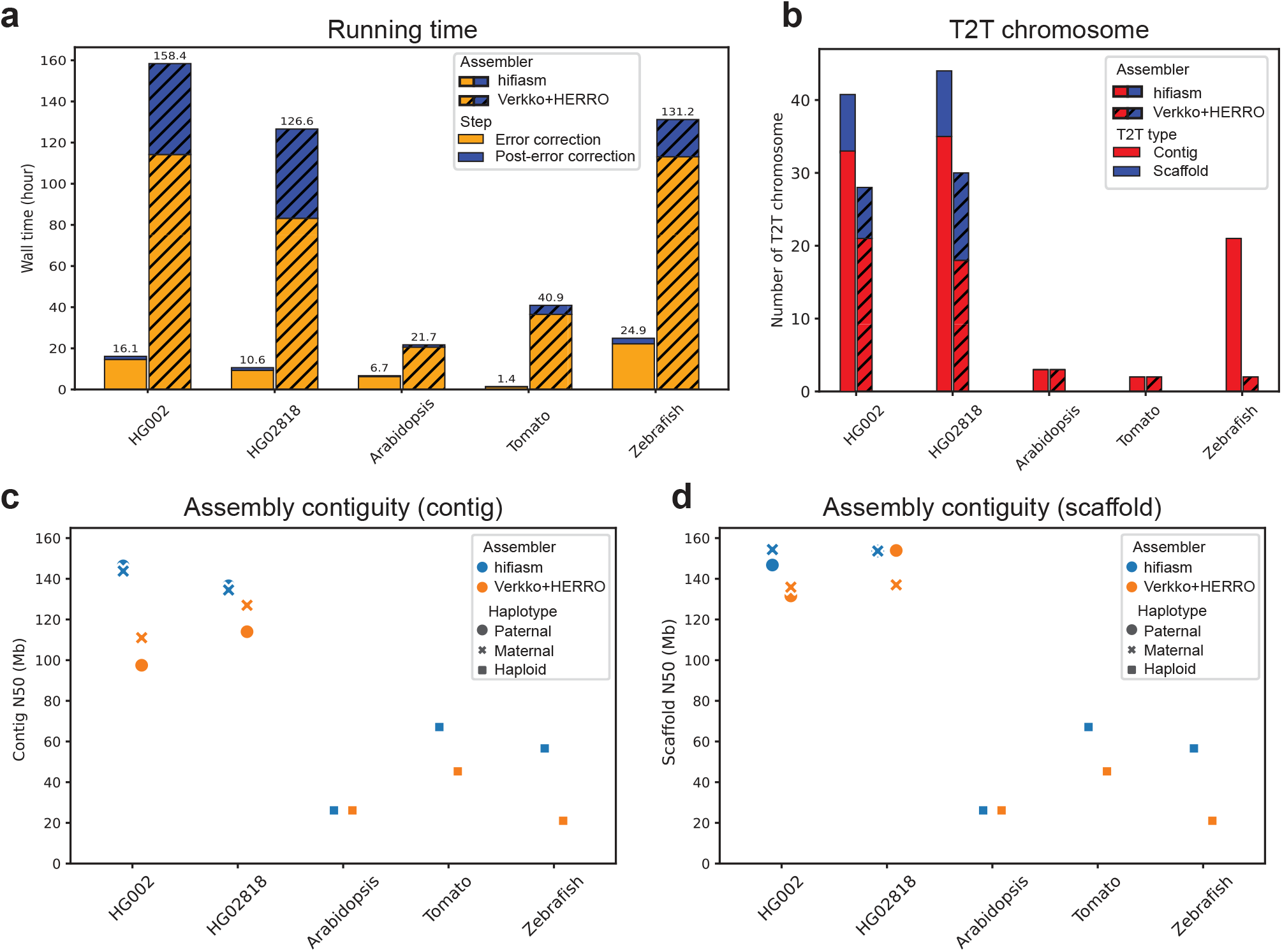
Assembly results using ONT ultra-long reads. All assemblies generated by hifiasm (ONT) and Verkko+HERRO are used as-is, except for the Arabidopsis assembly by hifiasm (ONT), in which low-coverage contigs were filtered out (Supplementary Section 1.1). **(a)** Running time of hifiasm (ONT) and Verkko+HERRO. Error correction and post-error correction times are shown separately for each sample. **(b)** Number of T2T contigs and scaffolds. T2T contigs and scaffolds indicate entire chromosomes reconstructed without and with gaps, respectively. The genomes of human (diploid), Arabidopsis (haploid), tomato (haploid), and zebrafish (haploid) have 46, 5, 12, and 25 chromosomes, respectively. **(c)** Contig N50. Assembly results for diploid human genomes (HG002 and HG02818) include two haplotypes, while the remaining three non-human genomes are haploid and have only one assembly. **(d)** Scaffold N50.

For HG002, using ultra-long reads instead of standard Simplex reads further improved assembly quality, as demonstrated by increased N50 values and a higher number of T2T contigs/scaffolds (see Fig. 2 and Table 1). For example, the number of T2T contigs produced by hifiasm (ONT) increased from 22 with standard reads to 33 with ultra-long reads. This ultra-long–based HG002 assembly of hifiasm (ONT) also reconstructs more T2T contigs and scaffolds than the recently reported Verkko HG002 assembly^8^requiring both HiFi and ultra-long reads. For the trio-binning assembly of HG002, hifiasm (ONT) produced 33 T2T chromosomes at the contig level and 44 at the scaffold level, while Verkko produced 22 T2T contigs and 32 T2T scaffolds combining both ONT ultra-long and PacBio HiFi reads.

### High-resolution reconstruction of challenging medically relevant genes

In clinical genomics, a key application of de novo assembly is the accurate reconstruction of challenging medically relevant genes that are often difficult to resolve using conventional alignment-based approaches. The GIAB Consortium has previously utilized hifiasm with PacBio HiFi reads to assemble a curated set of 273 medically important genes that are particularly difficult to reconstruct due to their high repetitiveness^36^. Despite the success of this strategy, more than 100 additional medically relevant genes remain unresolved in HiFi-based assemblies.

A representative example is the pair of highly homologous genes *SMN1* and *SMN2*. Biallelic pathogenic variants in *SMN1* lead to spinal muscular atrophy (SMA), a progressive neurodegenerative disorder characterized by muscle weakness and atrophy resulting from the loss of motor neurons in the spinal cord^37^. Accurately determining the sequence of *SMN1* and its paralog *SMN2* is critical for diagnosing SMA and informing therapeutic decisions^36, 38^. However, the HiFi-based assemblies used in the GIAB benchmarking framework are unable to fully resolve both *SMN1* and *SMN2*.

We evaluated the resolution of *SMN1* and *SMN2* in the haplotype-resolved assemblies generated by hifiasm (ONT) and Verkko+HERRO using both ONT standard and ultra-long reads. To minimize potential inaccuracies caused by reference bias, we aligned the HG002 assemblies to the HG002 Q100 reference^28^, which was generated from the same individual. Based on the annotated coordinates of *SMN1* and *SMN2*, we selected the 77.5–78.5 Mb region on *chr5 PATERNAL* and the 71.5–73 Mb region on *chr5 MATERNAL* of the HG002 reference as the target loci. We then aligned these regions to the corresponding paternal and maternal assemblies produced by each assembler to assess the accuracy and completeness of *SMN1* and *SMN2* reconstruction.

As shown in Fig. 3, hifiasm (ONT) successfully reconstructs both *SMN1* and *SMN2* using either ONT standard or ultra-long Simplex reads. Notably, this demonstrates that hifiasm (ONT) can resolve one of the most challenging medically relevant loci in the human genome using cost-effective and widely accessible ONT standard Simplex reads. This capability broadens the feasibility of T2T assembly for clinical samples, where ultra-long sequencing is often impractical. In contrast, Verkko+HERRO fails to faithfully reconstruct these regions. With ONT standard reads, it is unable to fully reconstruct either haplotype. When using ultra-long reads, Verkko+HERRO successfully assembles the paternal haplotype (Fig. 3c), but still fails to resolve the maternal haplotype (Fig. 3d). Hifiasm (ONT) outperforms Verkko+HERRO’s ultra-long read assemblies, even when using only ONT standard reads.

**Figure 3.**
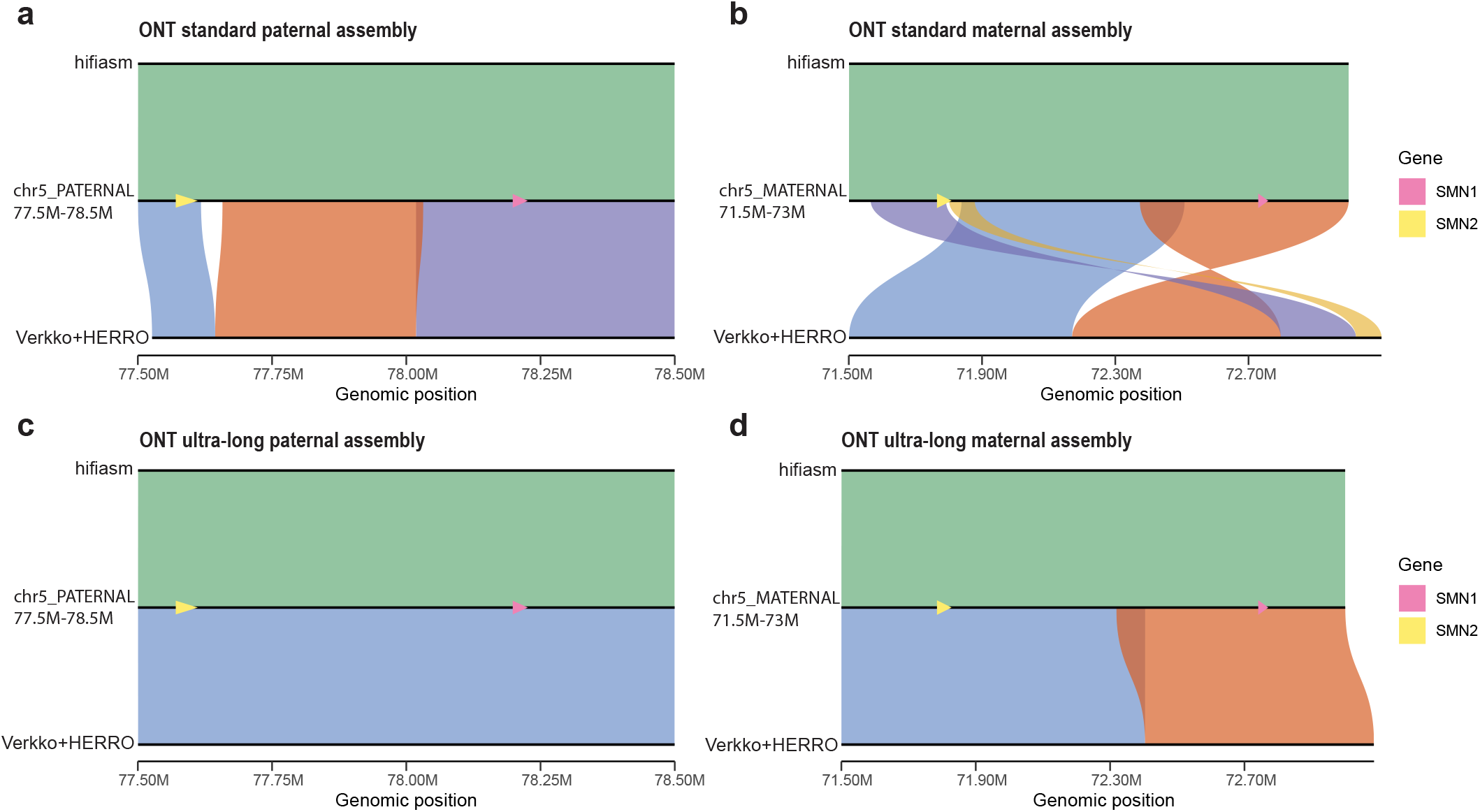
Comparison of HG002 assemblies with the HG002 T2T Q100 reference across the *SMN1/2* region. Each plot, generated using SVbyEye^31^, shows minimap2^22^ alignment results of the assemblies to the annotated *SMN1* and *SMN2* regions within the HG002 Q100 reference. Alignments corresponding to the same contig are shown in the same color. For consistency, several contigs aligned to the reverse-complement strand—such as the orange contig in Fig. 3b—are shown in their original orientation. Alignments of hifiasm (ONT) are shown above the reference, and alignments of Verkko+HERRO are shown below. The positions of *SMN1* and *SMN2* in the HG002 Q100 reference are highlighted in pink and yellow, respectively. **(a)** HG002 paternal assemblies using ONT standard Simplex reads. **(b)** HG002 maternal assemblies using ONT standard Simplex reads. **(c)** HG002 paternal assemblies using ONT ultra-long Simplex reads. **(d)** HG002 maternal assemblies using ONT ultra-long Simplex reads.

We then evaluated gene-level resolution in the fully phased HG002 assemblies. As in the multicopy gene retention analysis in Table 1, genes were aligned to each assembly, and resolution was assessed by comparing the aligned genes to those in the reference genome. Instead of using a generic reference as Table 1, here we used the HG002 Q100 reference from the same individual, ensuring an identical gene set across assemblies–critical for accurately evaluating highly repetitive or multi-allelic genes. As shown in Supplementary Table 7, the HiFi assemblies contained hundreds of unresolved genes, many within medically relevant and difficult-to-assemble loci. In contrast, the ONT assemblies generated by hifiasm (ONT) reduced the number of unresolved genes by an order of magnitude and achieved gene-level resolution comparable to ONT ultra-long assemblies even when using ONT standard reads. This is notable, as ONT standard reads are readily obtainable from clinical samples, whereas ONT ultra-long reads typically cannot. With ONT standard reads, Verkko+HERRO achieved lower gene-level resolution than hifiasm (ONT). Interestingly, the ONT standard-read assembly using Verkko+HERRO resolved more challenging genes than the HiFi assembly despite a lower contig N50, highlighting the importance of read length in resolving challenging medically relevant genes.

### Genome-wide patterns of assembly correctness in human genomes

To evaluate different assembly approaches, we again utilized alignment to the HG002 Q100 reference^28^. As shown in Fig. 4a, hifiasm (ONT) achieved the best results with ONT reads, outperforming the PacBio HiFi assembly and Verkko+HERRO assemblies. With hifiasm (ONT), ONT ultra-long reads not only yielded more contiguous assemblies than ONT standard reads but also resulted in the fewest misassemblies larger than 50 bp. Smaller misassemblies were ignored, as they can typically be resolved during post-assembly polishing. For samples lacking reference genomes, we employed Flagger^33^ and NucFlag^39^to perform reference-free evaluations by aligning reads back to their respective assemblies (Fig. 4b, c). The results again showed that hifiasm (ONT) generated the most accurate assemblies and that ONT assemblies were generally more accurate than PacBio HiFi assemblies. We also observed that different misassembly evaluation methods yielded notably different results (Supplementary Fig. 5), with reference-free approaches tending to overestimate misassemblies relative to alignment-based ground truths. This suggests that reliable misassembly evaluation remains a challenging problem. For HG002 fully phased assemblies, the genome-wide distribution of misassemblies was also assessed (Fig. 4d, e and Supplementary Fig. 4). The analysis revealed that most assembly errors occurred in highly repetitive regions such as centromeres.

**Figure 4.**
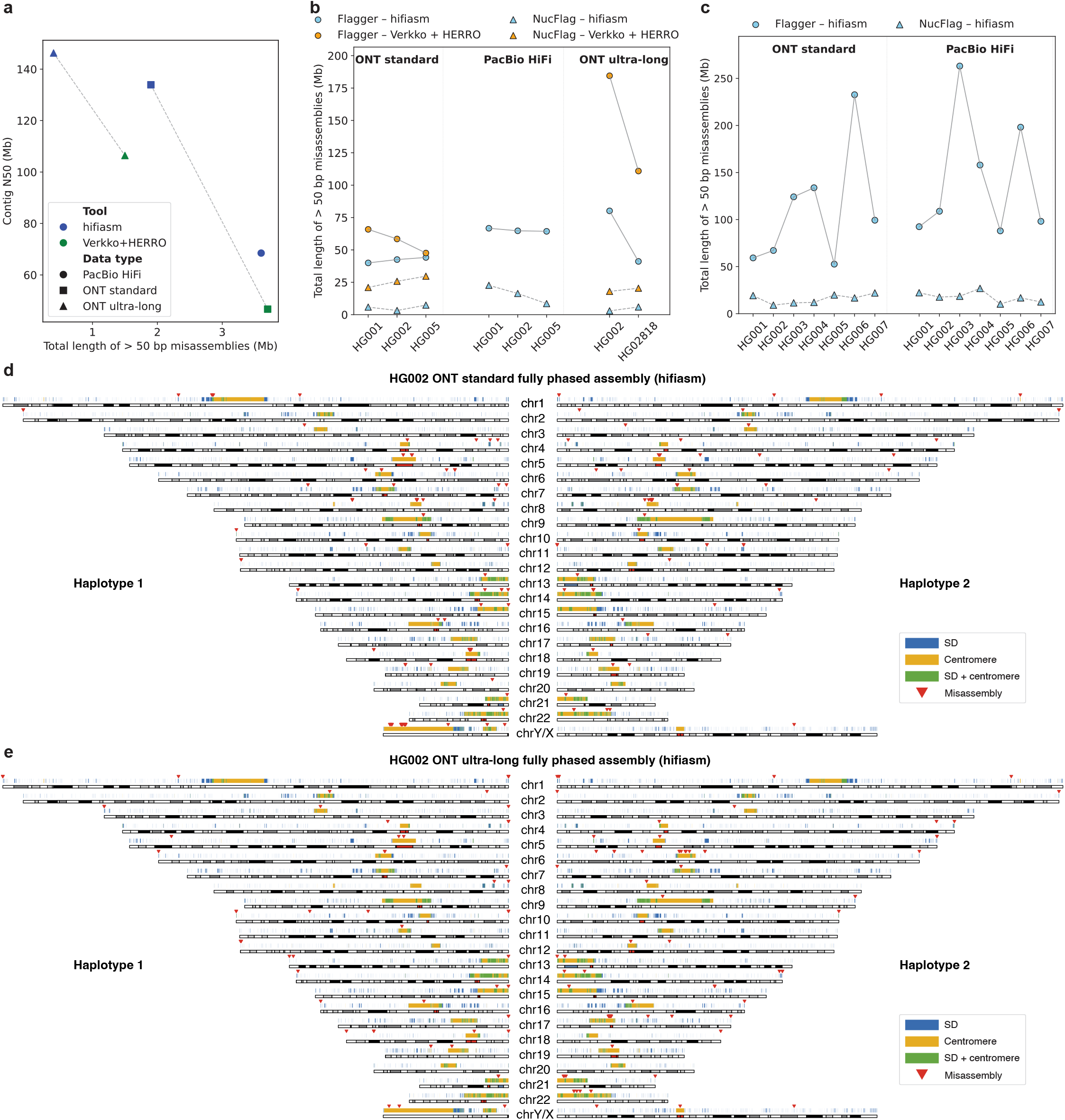
Evaluation of misassemblies in human genome assemblies. Only misassemblies larger than 50 bp are counted, as smaller errors can typically be corrected by polishing. For HG002-specific results (a), (d), and (e), misassemblies are accurately assessed against the HG002 Q100 ground truth. In contrast, other assemblies are approximately evaluated using Flagger^33^ and NucFlag^39^, as no ground truth is available. In panels (d) and (e), segmental duplication and centromere annotations are shown in blue and yellow, with overlapping regions highlighted in green and misassembly sites marked by red triangles. **(a)** Relationship between misassembly length and assembly contiguity in fully phased HG002 assemblies. **(b)** Misassemblies in all trio-binning human assemblies. **(c)** Misassemblies in all non-trio-binning human assemblies. **(d)** Genome-wide distribution of misassemblies in the HG002 ONT standard fully phased assembly produced by hifiasm (ONT). **(e)** Genome-wide distribution of misassemblies in the HG002 ONT ultra-long fully phased assembly produced by hifiasm (ONT).

### Characterization of unresolved assembly gaps in human genomes

We next analyzed the sequence composition of gaps that could not be resolved by the hifiasm toolkit. Assembly gaps were identified from alignments of contig ends that do not coincide with chromosome ends. For HG002, all contigs were mapped to the HG002 Q100 reference genome As shown in Fig. 5a–c, compared with the PacBio HiFi assembly, ONT assemblies resolved substantially more regions with high GA/TC content, low complexity, extreme GC/AT composition— regions that have traditionally been difficult to assemble using other sequencing technologies^6^. Ultra-long assemblies further resolved a substantially higher number of satellites, primarily from centromeric regions, indicating that ONT ultra-long reads remain essential for complete assembly of the human genome.

**Figure 5.**
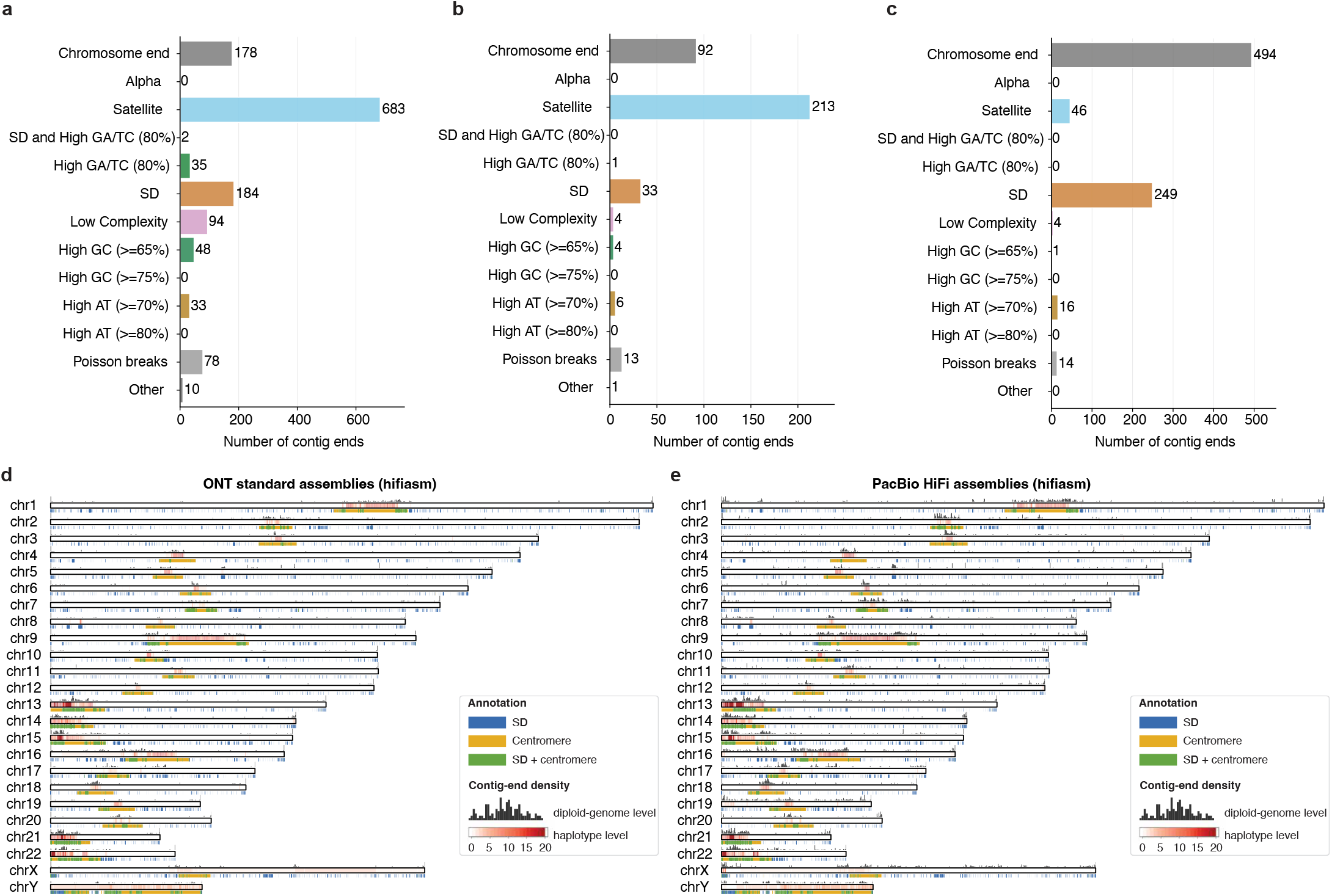
Genome-wide distribution of contig ends representing potential assembly gaps in human genome assemblies generated by hifiasm. Contig ends correspond either to chromosome ends or to unresolved assembly gaps. We aligned contig ends from each assembly to either the HG002 or CHM13 T2T reference genome. For the HG002-specific results **(a–c)**, evaluations were performed using the HG002 Q100 ground truth. For results (d) and (e), assemblies were aligned to the CHM13 T2T reference genome. (a–c) Number of contig ends in the HG002 phased assemblies produced by hifiasm using (a) PacBio HiFi, (b) ONT standard, and (c) ONT ultra-long reads. Each contig end was hierarchically classified according to its overlap with annotated sequence features. Poisson breaks refer to isolated contig ends (*≤*2 ends within a *±*100 kb window), which likely represent random assembly breaks. **(d–e)** Genome-wide distribution of assembly gaps across 10 diploid human genome assemblies (20 haplotypes) generated using (d) ONT standard Simplex and (e) PacBio HiFi reads, including both trio and non-trio assemblies. Chromosomal bars show haplotype-level gap coverage as a red heatmap, indicating the number of haplotypes with gaps at each position (max: 20 for autosomes, sex-aware for chrX/Y). Black bars above represent the number of diploid genomes with assembly gaps at each position (max: 10, after diploid-genome-level deduplication). Annotations below indicate segmental duplications (blue), centromeric satellites (yellow), and overlaps (green). More details are provided in Supplementary Section 1.13 and 1.14.

We also observed that ONT ultra-long assemblies contained more contig ends aligned to chromosome ends and segmental duplication regions. Most of these originated from small, erroneous contigs shorter than 500 kb, constructed from a limited number of reads with high sequencing error rates. By filtering out these small contigs, the number of contig ends within chromosome ends and segmental duplications was substantially reduced without compromising overall assembly quality, as these genomic regions were already represented in longer, higher-quality contigs. We hypothesize that the ONT standard assemblies contained fewer such erroneous contigs because ONT standard reads were processed using a newer version of the Dorado basecaller (v0.7.2), whereas the ONT ultra-long assemblies were generated from reads processed using an older version (v0.4.0). As Dorado continues to improve, base-level read accuracy will increase, reducing the frequency of erroneous short contigs.

Finally, we compared assembly gaps observed in PacBio HiFi and ONT Simplex standard human genome assemblies (Fig. 5d,e; 20 haplotypes in total for different assembly types, including both fully and partially phased assemblies). Because no complete reference genome is available for most samples, contigs were aligned to the CHM13 T2T reference. Although accurate alignment of contig ends can be challenging due to their repetitive and multi-allelic nature across individuals, we observed strong concordance: most assembly gaps clustered within repetitive regions. This observation is consistent with previous findings^6^and the distribution of misassemblies in Fig. 4d,e and Supplementary Fig. 4. Notably, ONT standard assemblies now resolve many highly challenging genomic regions that remain unresolved in PacBio HiFi assemblies (Fig. 5d,e). Based on haplotype-level density (red chromosomal bars), PacBio HiFi assemblies exhibit substantially more assembly gaps near chromosome ends (e.g., chr16, chr19, and chrX). However, because assembly gaps were measured by aligning assemblies to the CHM13 reference, there is alignment noise and uncertainty, making the haplotype-level density less accurate than the HG002-specific evaluation shown in Fig. 5a–c. This is largely because over 5% of each assembly could not be aligned to the CHM13 reference, and vice versa, with most assembly gaps located around these regions. Such regions are not only difficult to assemble^23^, often containing unresolved gaps, but also difficult to align^40^due to their high sequence divergence among individuals. To mitigate this, we also provide the diploid-genome level density (black bars above the chromosomal tracks), which represents the number of diploid genomes (out of 10 total) with at least one haplotype exhibiting contig ends at each position. Both haplotypes of each diploid genome were merged and counted as a single observation to prevent double-counting. This analysis more clearly demonstrates that ONT assemblies outperform PacBio HiFi assemblies across the genome.

## Discussion

Since the release of the first T2T human genome^23^, reconstructing entire genomes from telomere to telomere has become feasible and is attracting increasing attention. Nevertheless, achieving this goal remains challenging for both sequencing technologies and computational methods. We present hifiasm (ONT), an efficient de novo assembly algorithm designed to leverage the longer read lengths of ONT Simplex data to achieve T2T assemblies without relying on complex hybrid assembly strategies. It introduces an improved error correction approach that addresses the recurrent sequencing errors within ONT Simplex reads. This enables high-accuracy de novo assembly methods, originally developed for PacBio HiFi data, to be directly applied to ONT R10.4.1 Simplex reads. In contrast, existing tools—including current T2T hybrid assemblers—remain unable to perform de novo assembly of ONT Simplex reads as effectively as PacBio HiFi data. By fully utilizing the potential of ONT Simplex sequencing, hifiasm (ONT) achieves superior performance over other assemblers that fail to make effective use of this data type.

In comparison to HERRO, which was developed in parallel for error correction of ONT Simplex reads, hifiasm (ONT) is substantially faster due to its use of an efficient dynamic programming method instead of a time-consuming deep learning approach. This reduces the total assembly time by an order of magnitude and eliminates the need for multiple high-end GPUs (e.g., four A100, L40s, or RTX8000 GPUs for assembling a human genome), effectively removing a major computational barrier for ONT Simplex assembly. Furthermore, by tightly integrating and optimizing error correction with its ONT-specific assembly strategy, hifiasm (ONT) outperforms Verkko+HERRO by a wide margin across nearly all evaluation metrics.

With ONT ultra-long Simplex reads, hifiasm (ONT) reconstructs more human chromosomes from telomere to telomere than any other approach. It not only outperforms Verkko+HERRO using the same ultra-long data but also exceeds the performance of hybrid assemblies that combine ONT ultra-long and PacBio HiFi reads. Hifiasm (ONT) is also the first method to demonstrate that T2T genome assembly can be achieved using cost-effective and easily accessible ONT standard Simplex reads. This advancement enables population-scale T2T assembly and makes it feasible for clinical samples where ultra-long sequencing is often impractical. For example, hifiasm (ONT) is able to fully resolve the highly homologous and medically important gene pair *SMN1* and *SMN2*, which has remained unresolved in previous PacBio HiFi-based assemblies. We anticipate that hifiasm (ONT) will soon make routine near T2T genome assembly possible across various research and clinical applications.

## Methods

### Overview of hifiasm (ONT)

The existing hifiasm assembly toolkit consists of three approaches: the original hifiasm^2^, hifiasm (Hi-C)^5^, and hifiasm (UL)^9^, each designed for specific purposes. These methods are proposed respectively for trio-binning haplotype-resolved assembly using parental data, single-sample haplotype-resolved assembly using Hi-C reads, and T2T hybrid assembly. The core component shared by these methods is constructing a high-quality assembly graph through de novo assembly of PacBio HiFi reads. However, due to the limited length of PacBio HiFi reads, long repetitive genomic regions often cannot be fully resolved within this core assembly graph.

To address this limitation, longer but less accurate ONT Simplex reads, especially ultra-long reads, have been utilized by existing hybrid T2T assemblers such as hifiasm (UL)^9^and Verkko^7, 8^. However, these assemblers do not fully leverage ONT ultra-long reads because they cannot perform de novo assembly with them directly. The higher error rate and recurrent sequencing errors of ONT reads pose challenges for distinguishing genomic variations from sequencing errors. As a result, both hifiasm (UL) and Verkko use a two-stage strategy: first constructing an assembly graph from PacBio HiFi reads with de novo assembly, then aligning ONT ultra-long reads to resolve regions unresolved by HiFi reads alone. For ONT reads, this alignment-based approach remains inherently limited by inaccuracies and biases introduced in the HiFi-based assembly graph.

Our hifiasm (ONT) approach enhances this core capability by enabling effective de novo assembly of ONT Simplex reads, fully utilizing their longer length. Compared to HiFi-based assembly graphs, ONT-based assembly graphs tend to be more accurate, cleaner, and resolve more repetitive regions. This improved ONT-based assembly graph substantially improves existing methods in hifiasm toolkit such as trio-binning or Hi-C phased assembly. The specific strategies of hifiasm (ONT) for utilizing ONT Simplex reads in T2T assembly are detailed below.

### Error correction of ONT Simplex reads

Error correction generates near error-free reads by fixing sequencing errors in raw data, which is a critical step for genome assembly. To correct a given target read *R*, assembly algorithms need to first collect all related reads originating from the same genomic region. Traditional algorithms assume that any overlapping read belongs to the same genomic region as *R* if they share high sequence similarity. However, this approach fails to distinguish highly similar repeat copies or haplotypes. As a result, many false-positive reads from other repeats or haplotypes may be incorrectly used to correct *R*, leading to an overcorrection issue that may collapse repeats and haplotypes in the final assembly.

To address this problem when assembling PacBio HiFi reads, hifiasm employs a key assumption that most sequencing errors in PacBio HiFi data occur randomly and typically appear in only a single read. Fig. 1(a) shows an example. When correcting the target read *R*, hifiasm identifies mismatches through pairwise alignments between *R* and all overlapping reads. Mismatches supported by multiple overlaps are considered as informative sites representing true genomic variants, while those appearing in only one read are treated as sequencing errors and ignored. This strategy is essentially similar to widely used variant-calling methods. Subsequently, only overlapping reads showing no differences at these informative sites compared to *R* are utilized for correction.

However, this existing hifiasm error correction strategy is unsuitable for ONT Simplex reads (Fig. 1(b)). Although the overall sequencing error rate of ONT Simplex reads is not significantly higher than that of PacBio HiFi reads, ONT Simplex reads exhibit a higher frequency of recurrent, non-random errors. Therefore, the current hifiasm method would mistakenly identify these recurrent errors as informative sites. For a target Simplex read *R*, overlapping reads originating from the same genomic region would thus be incorrectly discarded if they differ from *R* at these recurrent error sites. As a result, most ONT Simplex reads would remain uncorrected, which cannot be used for de novo genome assembly.

In hifiasm (ONT), we introduce an approach that leverages the long-range phasing information of ONT reads to improve error correction. The basic idea is that true informative sites representing real genomic variants usually appear together and are mutually compatible with other informative sites. In practice, hifiasm (ONT) clusters potential informative sites based on their compatibility. As shown in Fig. 1(c), given a target ONT Simplex read awaiting correction, overlapping reads at a potential informative site (*x, y, m, n, z*, or *t*) can be classified into two phases: phase 0 (matching the target read) and phase 1 (differing from the target read). For two sites, if both consistently classify overlapping reads into identical phases, they are considered compatible and grouped together. True variants are expected to be compatible with multiple other real variant sites. In contrast, isolated sites lacking compatibility with others (such as *z* and *t*) have a higher likelihood of representing sequencing errors. This method is conceptually similar to leveraging haplotype phasing to improve the accuracy of variant calling. To further enhance reliability, hifiasm (ONT) adopts the following criteria: (i) An isolated site is considered informative only if it is supported by a sufficiently high number of reads; (ii) Any grouped site that is supported by more than one read is regarded as an informative site, as such sites are inherently more reliable.

In practice, it is necessary to develop an efficient algorithm for grouping potential informative sites, as this operation must be performed for each read during error correction. To achieve this, we propose a dynamic programming method designed to identify the largest compatible group for each site. Specifically, let *R* be the target read awaiting correction, and *S* be the list of *N* potential informative sites within *R*, sorted by their positions. Here, *S*[*i*] denotes the *i*-th site in the list, and *S*[*i*][*k*] represents the phase assignment of the *k*-th read at site *S*[*i*]. The value of *S*[*i*][*k*] can take one of the following states:

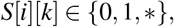

where 0 and 1 indicate that the *k*-th read is assigned to phase 0 or phase 1, respectively, and * indicates that the *k*-th read does not cover site *S*[*i*]. The details of the dynamic programming method are described as follows:

1. *Subproblem*. Let *LCG*[*i*] be the size of the largest compatible group in *S* that ends at index *i* and is compatible with *S*[*i*]. The goal is to compute *LCG*[0] to *LCG*[*N* −1] for all sites and identify those with values greater than 1, which indicate a compatible group rather than an isolated site.
2. *Recurrence relation*. Formally, the recurrence relation is defined as follows:

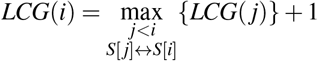

where *S*[ *j*] ↔ *S*[*i*] indicates that *S*[ *j*] is compatible with *S*[*i*]. Two sites *S*[ *j*] and *S*[*i*] are considered compatible if and only if

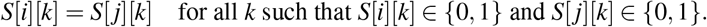

Fig. 1(c) illustrates an example of the dynamic programming matrix *LCG*.
3. *Traceback for grouping sites*. Hifiasm (ONT) identifies any entry where *LCG*[*i*] *>* 1, starting from the highest value and proceeding downward. For each site *S*[*i*] with *LCG*[*i*] *>* 1 that has not yet been assigned to a cluster, the algorithm traces back through its compatible prefix sites *S*[ *j*], following the path used to compute *LCG*(*i*). For example, in Fig. 1(c), hifiasm (ONT) starts from *LCG*(5), which holds the highest score, and groups the corresponding sites *S*[5], *S*[4], *S*[2], and *S*[0] (i.e., *n, m, y*, and *x*) during the traceback process.

The time and space complexity of this dynamic programming method are *O*(*n*^2^) and *O*(*n*), respectively, making it efficient for error correction in de novo assembly. An additional advantage of this approach is that it does not rely on the diploid genome assumption, enabling it to handle polyploid genomes or highly similar repeats with more than two repeat copies. As demonstrated in Fig. 1(d), hifiasm (ONT) successfully identifies *x* and *m* as one group and *n* and *y* as another group when there are three haplotypes available.

We further improve the error correction by filtering out low-quality base pairs as follows:

1. *Potential homopolymer sequencing errors*. ONT reads are known to exhibit a high sequencing error rate within homopolymer regions. If a potential informative site is located in a homopolymer region, hifiasm (ONT) discards it, as it is more likely to result from homopolymer-induced sequencing errors.
2. *Strand bias*. Given the target read *R*, hifiasm (ONT) excludes an informative site if all reads supporting *R* originate from one strand, while all other reads that differ from *R* at this site originate from the opposite strand. Strand bias is a common sequencing error observed in ONT reads.
3. *Low base quality score*. In addition to sequence data, hifiasm (ONT) also loads base quality scores into memory. Any base pair with a quality score below 10 is considered a potential sequencing error and excluded from the calculation of informative sites.

One potential challenge for hifiasm (ONT) arises when sequencing errors occur exactly at the position of a true variant, making that variant incompatible with others. However, in practice, such cases are rare. Even if a true variant in one read is affected by sequencing errors, other nearby variants that remain unaffected can still be accurately detected, allowing hifiasm (ONT) to effectively separate haplotypes and resolve repeat copies.

### Improved strategies for T2T assembly

With the error correction approach in hifiasm (ONT), most sequencing errors within ONT Simplex reads can be effectively corrected. These nearly error-free reads are then used with the existing assembly strategies in hifiasm to construct a high-quality assembly graph. For haploid genomes, a graph cleaning strategy is applied to produce linear assembly results. For diploid genomes, additional data—such as parental or Hi-C reads—are required to generate haplotype-resolved assemblies using hifiasm’s existing trio-binning or Hi-C phasing approaches.

To further improve T2T assembly, we developed a strategy to retain telomere sequences in the final assembly. A common issue in hifiasm is that while it can reconstruct entire chromosomes, it may still miss telomeric sequences at chromosome ends. This occurs because, in the assembly graph—particularly for diploid genomes—telomere ends often appear as tips. During graph cleaning, hifiasm typically discards these tips, as most are caused by assembly errors. As a result, telomere sequences may be inadvertently removed from the final assembly. To address this, the improved T2T assembly strategy in hifiasm (ONT) first checks whether any reads contain telomeric sequences before the assembly. If such reads are detected, hifiasm preserves the corresponding graph tips during graph cleaning. This approach helps retain more telomere ends and results in an increased number of T2T contigs and scaffolds.

We also developed a dual-scaffold approach to assemble more chromosomes from telomere to telomere at the scaffold level. The goal is to scaffold gapless contigs into longer, gapped scaffolds by leveraging information from both haplotypes. The basic idea is that, for an assembly gap in haplotype 1, the dual-scaffold approach examines the corresponding homologous regions in haplotype 2. If the region in haplotype 2 is completely assembled without gaps, the dual-scaffold method fills the gap in haplotype 1 with Ns, using the estimated length inferred from the complete sequence in haplotype 2. In essence, this approach performs reference-guided scaffolding for each haplotype^41^, using the other haplotype as a reference.

### Library preparation, sequencing, and basecalling

ONT standard Simplex sequencing data for the GIAB samples HG001–HG007 have been deposited in the official Oxford Nanopore Technologies open data repository (s3://ont-ope n-data/giab 2025.01/). Cell lines for these samples were obtained from the Human Genetic Cell Repository at the Coriell Institute for Medical Research and cultured according to the supplier’s recommended protocols. High-molecular-weight DNA was extracted using the QIAgen Puregene cell extraction kit, followed by library preparation with the SQK-LSK114 kit according to ONT protocols, and sequencing was performed on PromethION flow cells using P48 instruments. Basecalling was conducted using Dorado v0.7.2 with both HAC v5.0.0 and SUP v5.0.0 models. For HG001, HG003, HG004, HG005, HG006, and HG007, reads from two flow cells were basecalled and used for assembly. For HG002, only data from a single flow cell was used, as one flow cell was sufficient to produce ONT Simplex reads at approximately 50× coverage. In addition, we re-basecalled the existing *Danio rerio* (zebrafish) dataset using Dorado v0.8.3 with the SUP v5.0.0 model to improve read-level base accuracy.

## Acknowledgements

This study was supported by the U.S. National Institutes of Health (grants R00HG012798 and RM1HG011558 to H.C.; R01HG010040, U01HG010971, and U41HG010972 to H.L.; and UM1DA058230 and R01CA269805 to P.J.P.). We thank the Human Pangenome Reference Consortium for making ultra-long datasets publicly available.

## Author contributions

H.C. designed the algorithm, implemented hifiasm (ONT), and drafted the manuscript. H.C. benchmarked hifiasm (ONT). H.Q. designed the evaluation of genome assemblies and led the comprehensive benchmarking of all approaches, supervised by P.J.P. S.M., K.R.L., R.W., and M.V. collected nanopore sequencing data and contributed to the assembly evaluation. H.L. directed the entire project. All authors contributed to writing the manuscript.

## Data availability

Human reference genome: GRCh38, CHM13v2, and HG002 Q100; ONT standard reads of HG001 (SUP): s3://ont-open-data/ giab 2025.01/basecalling/sup/HG001; ONT standard reads of HG001 (HAC): s3://ont-open-data/giab 2025.01/basecalling /hac/HG001; PacBio HiFi reads of HG001: https://ftp.ncbi.nlm.nih.gov/ReferenceSamples/giab/data/NA12878/HudsonAl_pha_PacBio_CCS/; Illumina short reads of HG001 and parents: ERR194147 (HG001), ERR194160 (paternal), ERR194161 (maternal); ONT standard reads of HG002 (SUP): s3://ont-open-data/giab 2025.01/basecalling/sup/HG002/PAW70337; ONT standard reads of HG002 (HAC): s3://ont-open-data/giab 2025.01/basecalling/hac/HG002/PAW70337; ONT ultra-long reads of HG002: https://s3-us-west-2.amazonaws.com/human-pangenomics/index.html?prefix=submissions/5b73fa0e-658a-4248-b2b8-cd16155bc157--UCSC_GIAB_R1041_nanopore/HG002_R1041_UL/dorado/v0.4.0_wMods/*ULCIR*.bam and https://s3-us-west-2.amazonaws.com/human-pangenomics/index.html?prefix=submissions/5b73fa0e-658a-4248-b2b8-cd16155bc157--UCSC_GIAB_R1041_nanopore/HG002_R1041_UL/dorado/v0.4.0_wMods/*ULNEB*.bam; PacBio HiFi reads of HG002: https://ftp.ncbi.nlm.nih.gov/ReferenceSamples/giab/data/AshkenazimTrio/HG002_N_A24385_son/PacBio_HiFi-Revio_20231031/; Illumina short reads of HG002 and parents: https://s3-us-west-2.amazo_naws.com/human-pangenomics/index.html?prefix=working/HPRC_PLUS/HG002/raw_data/Illumina/: ONT standard reads of HG003 (SUP): s3://ont-open-data/giab 2025.01/basecalling/sup/HG003; ONT standard reads of HG003 (HAC): s3://ont-open-data/giab 2025.01/basecalling/hac/HG003; PacBio HiFi reads of HG003: https://ftp.ncbi.nlm.nih.gov/ReferenceSamples/giab/data/AshkenazimTrio/HG003_NA24149_father/PacBio_HiFi-Revio_20231031/ and https://ftp.ncbi.nlm.nih.gov/ReferenceSamples/giab/data/AshkenazimTrio/HG003_NA24149_father/PacBio_CCS_Google_15kb/; Illumina short reads of HG003: https://s3-us-west-2.amazonaws.com/human-pangenomics/index.html?prefix=working/HPRC_PLUS/HG002/raw_data/Illumina/parents/HG003/; ONT standard reads of HG004 (SUP): s3://ont-open-data/giab_2025.01/basecalling/sup/HG004; ONT standard reads of HG004 (HAC): s3://ont-open-data/giab_2025.01/basecalling/hac/HG004; PacBio HiFi reads of HG004: https://ftp.ncbi.nlm.nih.gov/ReferenceSamples/giab/data/AshkenazimTrio/HG004_NA24143_mother/PacBio_HiFi-Revio_20231031/ and https://ftp.ncbi.nlm.nih.gov/ReferenceSamples/giab/data/AshkenazimTrio/HG004_NA2_4143_mother/PacBio_CCS_Google_15kb/; Illumina short reads of HG004: https://s3-us-west-2.amazonaws.com/human-pangenomics/index.html?prefix=working/HPRC_PLUS/HG002/raw_data/Illumina/parents/HG004/; ONT standard reads of HG005 (SUP): s3://ont-open-data/giab 2025.01/basecalling/sup/HG005; ONT standard reads of HG005 (HAC): s3://ont-open-data/giab 2025.01/basecalling/hac/HG005; PacBio HiFi reads of HG005: https://ftp.ncbi.nlm.nih.gov/ReferenceSamples/giab/data/ChineseTrio/HG005_NA24631_son/PacBio_CCS_15kb_20kb_hemistry2/uBAMs/; Illumina short reads of HG005: https://s3-us-west-2.amazonaws.com/human-pangenomics/index.html?prefix=working/HPRC_PLUS/HG005/raw_data/Illumina/; ONT standard reads of HG006 (SUP): s3://ont-open-data/giab 2025.01/basecalling/sup/HG005; ONT standard reads of HG006 (HAC): s3://ont-open-data/giab 2025.01/basecalling/hac/HG005; PacBio HiFi reads of HG006: https://ftp.ncbi.nlm.nih.gov/ReferenceSamples/giab/data/ChineseTrio/HG006_NA24694-huCA017E_father/PacBio_CCS_15kb_20kb_chemistry2/uBAMs/ and https://ftp.ncbi.nlm.nih.gov/ReferenceSamples/giab/data/ChineseTrio/HG006_NA24694-huCA017E_father/PacBio_HiFi_Google/; Illumina short reads of HG006: https://s3-us-west-2.amazonaws.com/human-pangenomics/index.html?prefix=working/HPRC_PLUS/HG005/raw_data/Illumina/parents/HG006/; ONT standard reads of HG007 (SUP): s3://ont-open-data/giab 2025.01/basecalling/sup/HG007; ONT standard reads of HG007 (HAC): s3://ont-open-data/giab 2025.01/basecalling/hac/HG007; PacBio HiFi reads of HG007: https://ftp.ncbi.nlm.nih.gov/ReferenceSamples/giab/data/ChineseTrio/HG007_NA24695-hu38168_mother/PacBio_CCS_15kb_20kb_chemistry2/uBAMs/ and https://ftp.ncbi.nlm.nih.gov/ReferenceSamples/giab/data/ChineseTrio/HG007_NA24695-hu38168_mother/PacBio_HiFi_Google/; Illumina short reads of HG007: https://s3-us-west-2.amazonaws.com/human-pangenomics/index.html?prefix=working/HPRC_PLUS/HG005/raw_data/Illumina/parents/HG007/; ONT ultra-long reads of HG02818: https://s3-us-west-2.amazonaws.com/human-pangenomics/index.html?prefix=working/HPRC_PLUS/HG02818/raw_data/nanopore/dorado0.7.2_sup4.1.0_5mCG_5hmCG/; Illumina short reads of HG02818 and parents: https://s3-us-west-2.amazonaws.com/human-pangenomics/index.html?prefix=working/HPRC_PLUS/HG02818/raw_data/Illumina/; ONT ultra-long reads of *Arabidopsis thaliana*: SRR29061597; Illumina short reads of *Arabidopsis thaliana*: https://ngdc.cncb.ac.cn/gsa/browse/CRA005350; ONT ultra-long data of *Danio rerio*: https://genomeark.s3.amazonaws.com/index.html?prefix=species/Danio_rerio/fDanRer17/genomic_data/ont/pod5/ and https://genomeark.s3.amazonaws.com/index.html?prefix=species/Danio_rerio/fDanRer17/genomic_data/ont/fast5/; Illumina short reads of *Danio rerio*: https://www.ncbi.nlm.nih.gov/sra?linkname=bioproject_sra_all&from_uid=1029986; ONT ultra-long data of *Solanum lycopersicum*: https://obj.umiacs.umd.edu/marbl_publications/duplex/Solanum_lycopersicum_heinz1706/UL/R10.4_40x.noduplex.fastq.gz; Illumina short reads of *Solanum lycopersicum*: https://ngdc.cncb.ac.cn/gsa/browse/CRA003995/CRX232533; ONT reads of *Linum usitatissimum* (Flax): SRR31124331; ONT reads used for comparison with Napu (Shasta): https://s3-us-west-2.amazonaws.com/human-pangenomics/index.html?prefix=publications/Napu_paper_ONT_Coriell_SingleFC_2023/HG0*_R10/reads/*.bam; Napu (Shasta) assemblies: https://s3-us-west-2.amazonaws.com/human-pangenomics/index.html?prefix=publications/Napu_paper_ONT_Coriell_SingleFC_2023/HG0*_R10/assembly/*contigs*fasta; All evaluated assemblies generated by hifiasm and Verkko+HERRO are available at: https://zenodo.org/records/15203417 (human genome assemblies from ONT reads), https://zenodo.org/records/17613526 (non-human genome assemblies from ONT reads), and https://zenodo.org/records/15205178 (assemblies from PacBio HiFi reads). All links and accession identifiers for the sequencing data are also listed in Supplementary Table 8.

## Code availability

Hifiasm (ONT) is available at https://github.com/chhylp123/hifiasm.

## Reporting Summary

Further information on research design is available in the Nature Research Reporting Summary linked to this article

