## Supplementary Information for "Efficient near telomere-to-telomere assembly of Nanopore Simplex reads"

#### 1 Software commands

##### 1.1 Hifiasm

To produce partially phased assemblies from ONT reads for diploid human genomes, we ran `hifiasm` (version 0.25.0-r726) using the dual assembly model with the following command:

```
hifiasm -o <outputPrefix> -t <nThreads> --ont --dual-scaf --telo-m CCCTAA <ONT-reads.fastq>
```

For trio-binning assemblies, we first generated paternal and maternal *k*-mer indexes using `yak` (version 0.1-r62-dirty) with the following commands:

```
yak count -b37 -t <nThreads> -o <pat.yak> <paternal-short-reads.fastq>
yak count -b37 -t <nThreads> -o <mat.yak> <maternal-short-reads.fastq>
```

We then performed trio-binning assembly with:

```
hifiasm -o <outputPrefix> -t <nThreads> --ont -1 <pat.yak> -2 <mat.yak> --dual-scaf --telo-m CCCTAA
<ONT-reads.fastq>
```

To produce assemblies from PacBio HiFi reads, we used the same command lines as above, omitting the `--ont` option. For partially phased assemblies:

```
hifiasm -o <outputPrefix> -t <nThreads> --dual-scaf --telo-m CCCTAA <HiFi-reads.fastq>
```

For trio-binning with HiFi reads:

```
hifiasm -o <outputPrefix> -t <nThreads> -1 <pat.yak> -2 <mat.yak> --dual-scaf --telo-m CCCTAA
<HiFi-reads.fastq>
```

We also assembled haploid non-human genomes with ONT reads:

```
hifiasm -o <outputPrefix> --ont -t <nThreads> -l0 --telo-m <telomere-motif> <ONT-reads.fastq>
```

The telomere motifs were specified using the `--telo-m` option as follows: `TTTAGGG` for *Arabidopsis thaliana* (arabidopsis), `CCCTAA` for *Danio rerio* (zebrafish), and `TTTAGGG` for both *Solanum lycopersicum* (tomato) and *Linum usitatissimum* (flax). For *Arabidopsis thaliana*, all low-coverage contigs with coverage $\leq 90$ (as indicated by the `rd:i:` tag in the GFA file) were filtered out. Most of these contigs, based on BLAST results, are of bacterial origin and likely represent contamination.

##### 1.2 HERRO

To produce ONT-only assemblies, all Simplex reads were first error-corrected using `HERRO` (R10.4.1 model) and then assembled with `Verkko`. The `HERRO` error correction tool (https://github.com/lbcb-sci/herro) was executed using scripts from commit 075306685b7a16e7a5cce2ab370530f64850506e (https://github.com/lbcb-sci/herro/tree/075306685b7a16e7a5cce2ab370530f64850506e). For the inference step, the R10.4.1 model available at http://metals.zesoi.fer.hr:9080/herro/model_v0.1.pt was used. The `HERRO` pipeline consists of three main steps: read preprocessing, creation of self-alignments with batching, and error correction using a deep learning model. Specifically, read preprocessing was first performed with the following command:

```
$herro/scripts/preprocess.sh <input_fastq> <output_prefix> <number_of_threads> <parts_to_split_job_into>
seqkit seq -ni <preprocess_output.fastq> > <preprocess_output_readID.txt>
```

Self-alignments were then generated using `minimap2` via the following command:

```
$herro/scripts/create_batched_alignments.sh <preprocess_output.fastq> <preprocess_output_readID.txt>
<number_of_threads> <alignment_batch_directory>
```

Finally, corrected reads were generated using the following command:

```
herro inference --read-alns <directory_alignment_batches> -t <feat_gen_threads_per_device> -d <gpus>
-m <model_path> -b <batch_size> <preprocess_output.fastq> <fasta_output>
```

#### 1.3 Verkko

`Verkko` v2.2.1 was run using the `HERRO`-corrected reads. To fully leverage the power of ONT data, `Verkko` was executed in hybrid assembly mode, where the corrected ONT reads were treated as PacBio HiFi reads, and the raw ONT reads were used as ultra-long reads. Following `HERRO`'s recommendations and based on our benchmarking for optimal assembly quality, the corrected reads were downsampled to 35× coverage when the original coverage substantially exceeded this threshold (i.e., >39×). Downsampling of `HERRO`-corrected reads to a target coverage was performed with `Seqtk` (version 1.4-r130-dirty) using the following command:

```
seqtk sample <herro_corrected.fasta> <proportion> > <downsampled.fasta>
```

Coverage was calculated by dividing the total number of base pairs by the estimated genome size. The estimated genome sizes used were 3.1 Gb for human, 1.5 Gb for *Arabidopsis thaliana* (Arabidopsis), 782 Mb for *Danio rerio* (zebrafish), 135 Mb for *Solanum lycopersicum* (tomato), and 480 Mb for *Linum usitatissimum* (flax).

For the haploid genome assemblies of *Arabidopsis thaliana* (Arabidopsis), *Danio rerio* (zebrafish), *Solanum lycopersicum* (tomato), and *Linum usitatissimum* (flax), `Verkko` was run with the following command:

```
verkko -d <output_directory> --hifi <downsampled.fasta> --nano <ont_raw.fastq>
   --local-memory <ram> --local-cpus <num_cpus> --haploid
```

To produce trio-binning assemblies of the human genome, we first generated *k*-mer databases from the assembly sample and parental short reads using `Meryl` (version 1.4.1):

```
meryl count compress k=30 threads=<threads> <input.fastq.gz> output=<prefix_compressed_k30.meryl>
```

Next, haplotype-specific *k*-mers (hapmers) were generated for use in `Verkko` assembly using `Merqury` (version 1.3, commit `ed8c3ba3ea8897d9151a7da8772cd5d83fbce474`):

```
$MERQURY/trio/hapmers.sh <maternal_compressed_k30.meryl> <paternal_compressed_k30.meryl>
<child_compressed_k30.meryl>
```

Finally, human genomes were assembled using the `HERRO`-corrected reads, raw ONT reads, and the pre-built hapmers:

```
verkko -d <output_directory> --hifi <downsampled.fasta> --nano <ont_raw.fastq>
   --local-memory <ram> --local-cpus <num_cpus>
   --hap-kmers <maternal_compress_k30.hapmer.meryl> <paternal_compress_k30.hapmer.meryl> trio
```

#### 1.4 Computing environment

All hifiasm assemblies were run using 64 CPUs and 1 TB of RAM. As the combination of `Verkko` and `HERRO` requires significantly more computational resources—including both CPU and GPU—different steps were executed on different servers to optimize performance. The detailed computing resources used for the `HERRO`+`Verkko` pipeline are summarized as follows:

1. HG001 (standard reads):
   - Non-deep learning step of `HERRO`: 68 CPUs
   - Deep learning step of `HERRO`: 4 RTX8000 GPUs (each with 48 GB VRAM) and 10 CPUs
   - `Verkko`: 96 CPUs
2. HG002 (standard reads):
   - Non-deep learning step of `HERRO`: 40 CPUs
   - Deep learning step of `HERRO`: 4 L40s GPUs (each with 48 GB VRAM) and 10 CPUs
   - `Verkko`: 96 CPUs
3. HG005 (standard reads):
   - Non-deep learning step of `HERRO`: 68 CPUs
   - Deep learning step of `HERRO`: 4 L40s GPUs (each with 48 GB VRAM) and 10 CPUs
   - `Verkko`: 96 CPUs
4. HG02818 (ultra-long reads):

    - Non-deep learning step of `HERRO`: 68 CPUs
    - Deep learning step of `HERRO`: 4 A100 GPUs (each with 80 GB VRAM) and 10 CPUs
    - `Verkko`: 96 CPUs

5. HG002 (ultra-long reads):
    - Non-deep learning step of `HERRO`: 68 CPUs
    - Deep learning step of `HERRO`: 4 A100 GPUs (each with 80 GB VRAM) and 10 CPUs
    - `Verkko`: 96 CPUs
6. *Arabidopsis thaliana* (ultra-long reads):
    - Non-deep learning step of `HERRO`: 68 CPUs
    - Deep learning step of `HERRO`: 2 L40s GPUs (each with 48 GB VRAM) and 1 CPUs
    - `Verkko`: 96 CPUs
7. *Solanum lycopersicum* (ultra-long reads):
    - Non-deep learning step of `HERRO`: 68 CPUs
    - Deep learning step of `HERRO`: 2 L40s GPUs (each with 48 GB VRAM) and 1 CPUs
    - `Verkko`: 96 CPUs
8. *Danio rerio* (ultra-long reads):
    - Non-deep learning step of `HERRO`: 68 CPUs
    - Deep learning step of `HERRO`: 8 A100 GPUs (each with 48 GB VRAM) and 10 CPUs
    - `Verkko`: 96 CPUs
9. *Linum usitatissimum* (standard reads):
    - Non-deep learning step of `HERRO`: 30 CPUs
    - Deep learning step of `HERRO`: 2 V100 GPUs (each with 32 GB VRAM) and 15 CPUs
    - `Verkko`: 96 CPUs

#### 1.5 T2T assessment

The script from the Human Pangenome Reference Consortium (HPRC) (https://github.com/biomonika/HPP/blob/main/assembly/wdl/workflows/assessAsemblyCompletness.wdl) was employed to identify telomere-to-telomere (T2T) contigs and scaffolds. Telomere motifs were species-specific:

- **Human**: `TTAGGG`
- ***Danio rerio*** (zebrafish): `TTAGGG`
- ***Arabidopsis thaliana***: `TTTAGGG`
- ***Solanum lycopersicum*** (tomato): `TTTAGGG`
- ***Linum usitatissimum*** (flax): `TTTAGGG`

#### 1.6 Assembly assessment using yak

The `yak` toolkit (version 0.1-r62-dirty) was used to evaluate assembly quality through the following three-step process:

1. Creation of `yak` files from short reads:

```
yak count -b37 -t<num_threads> -o <output.yak> <(zcat sr*.fastq.gz)> <(zcat sr*.fastq.gz)>
```

2. Calculating assembly quality value (QV):

```
yak qv -t<num_threads> <child.yak> <assembly.fasta> > <assembly_qv.txt>
```

3. Calculating hamming and switch errors for haplotype evaluation:

```
yak trioeval -e -t<num_threads> <paternal.yak> <maternal.yak> <hap1_assembly.fasta>
> <hap1.trioeval.txt>
yak trioeval -e -t<num_threads> <paternal.yak> <maternal.yak> <hap2_assembly.fasta>
> <hap2.trioeval.txt>
```

#### 1.7 Assembly assessment using Merqury

Assembly quality was evaluated using `Merqury` (version 1.3, commit `ed8c3ba3ea8897d9151a7da8772cd5d83fbce474`) together with `Meryl` (version 1.4.1), following the workflow below:

For samples with trio data (e.g., diploid human genomes):

1. *k*-mer databases were constructed with `Meryl`:

```
meryl count k=<k> threads=<threads> memory=<memory> <*.fastq.gz> output=<output_prefix.meryl>
```

2. Haplotype-specific *k*-mers were generated:

```
$MERQURY/trio/hapmers.sh <maternal.meryl> <paternal.meryl> <child.meryl>
```

3. Assembly evaluation for diploid samples was performed:

```
$MERQURY/merqury.sh <child.meryl> <maternal.hapmer.meryl> <paternal.hapmer.meryl>
    <hap1.fasta> <hap2.fasta> <output_prefix>
```

4. Hamming and switch error rates were calculated:

```
$MERQURY/trio/hamming_error.sh <output_prefix.hapmers.count> <hap1.fasta> <hap2.fasta>
```

For samples without trio data (e.g., Arabidopsis, tomato, zebrafish, HG003, HG004, HG006, HG007):

1. *k*-mer databases were constructed from both short and HiFi reads:

```
meryl count k=<k> threads=<threads> memory=<memory> <*.short.fastq.gz> output=<short_prefix.meryl>
meryl count k=<k> threads=<threads> memory=<memory> <*.hifi.fastq.gz> output=<hifi_prefix.meryl>
```

2. A hybrid *k*-mer database was created by filtering and merging:

```
meryl greater-than 1 <short_prefix.meryl> output=<short_prefix.gt1.meryl>
meryl greater-than 1 <hifi_prefix.meryl> output=<hifi_prefix.gt1.meryl>
meryl union-sum <short_prefix.gt1.meryl> <hifi_prefix.gt1.meryl> output=<child.meryl>
```

3. Final assessment was performed using:

```
$MERQURY/merqury.sh <child.meryl> <assembly.fasta> <output_prefix>
```

*K*-values were selected based on recommendations from `Merqury`'s `best_k.sh` script:

- Human: 21
- *Danio rerio* (zebrafish): 20
- *Solanum lycopersicum* (tomato): 19
- *Arabidopsis thaliana*: 18

#### 1.8 Running asmgene

For all human genome assemblies (see Table 1 in the main manuscript), we evaluated gene completeness by aligning cDNAs to both the CHM13v2 reference genome and the assembled contigs using `minimap2` (version 2.28-r1209). Gene-level completeness was then assessed using `paftools.js`, a utility from the `minimap2` package. The commands used are as follows:

```
minimap2 -cxsplice:hq -t <nThreads> <ref.fa> <cDNAs.fa> > <ref.paf>
minimap2 -cxsplice:hq -t <nThreads> <asm.fa> <cDNAs.fa> > <asm.paf>
paftools.js asmgene -a -i.97 <ref.paf> <asm.paf>
```

For HG002-specific evaluation (see Supplementary Table 7), the HG002 Q100 reference was used instead of the CHM13v2 reference genome. Because the reference and assemblies originate from the same individual, we increased the sequence identity threshold between genes identified in the reference and the assembly to 99%:

```
paftools.js asmgene -a -i.99 <ref.paf> <asm.paf>
```

#### 1.9 SVbyEye

The alignment used for SVbyEye visualization was generated with the following command:

```
minimap2 -t <nThreads> --eqx --secondary=no -c -x asm20 asm_contig.fa hg002v1.1.SMN12.gene.fa
```

#### 1.10 Immuannot

Commit 31362c3 was used. Because Immuannot is based on gene sequence alignment, novel genes may not be detected if they differ substantially from the reference sequences. In addition, when processing assemblies containing a large number of contigs ($>$ 200), the default -N parameter in minimap2 can cause certain genes to be missed entirely rather than classified as present or absent. To address this issue, we implemented a ”split contigs” strategy in which each assembly was divided into multiple files, each containing fewer than 200 contigs. These files were processed separately by Immuannot, and the results were subsequently merged. This approach circumvents the -N parameter limitation and ensures comprehensive analysis of all target genes.

#### 1.11 Assembly-based variant calling

To evaluate assembly-based variant calling performance (see Supplementary Table 6), variant calls for each assembly were generated using Dipcall v0.3, and benchmarked against the GIAB HG002 T2TQ100-v1.0 truth set and corresponding confident regions on GRCh38. For Dipcall outputs, input VCFs were normalized prior to comparison: multiallelic sites were split, variants were left-aligned, deduplicated, and sorted/indexed using bcftools and tabix, with the same GRCh38 reference FASTA used by the truth set.

Benchmarking was performed using hap.py v0.3.15 with parameters --engine=vcfeval and --gender male, restricted to GIAB confident regions. To assess performance in sequence-context–specific settings, a two-class stratification table distinguishing homopolymer and non-homopolymer regions was used from the GIAB GRCh38 stratifications v3.6 (https://ftp-trace.ncbi.nlm.nih.gov/ReferenceSamples/giab/release/genome-stratifications/v3.6/GRCh38@all/LowComplexity/). For each sequencing condition, hap.py produced both genome-wide and per-stratum summaries. From these outputs, precision, recall, and F1 scores were calculated for SNPs, INDELs, and their union (“ALL”), considering only PASS variants.

#### 1.12 Misassembly detection

To comprehensively assess misassemblies, we employed three approaches: Flagger, NucFlag, and reference-based evaluation specifically for HG002. Only misassemblies larger than 50 bp were considered, as smaller discrepancies are more likely to reflect consensus errors in base-level rather than true structural misassemblies and can typically be corrected through polishing after assembly. The detailed procedures for each method are described below.

##### 1.12.1 Flagger-based detection

For Flagger v1.1.0, PacBio HiFi reads were aligned to the combined haplotype 1 and haplotype 2 diploid assembly using minimap2 v2.30-r1287 (-x map-hifi -k 19 --eqx --cs -Y -L). The Flagger Singularity pipeline was then executed as follows:

1. Create a whole-genome BED file:

```
# Go to the working directory
# Place FASTA, BAM, and BAM index files in this directory
cd ${WORKING_DIR}

# Create a FASTA index (assuming the file is not gzipped)
samtools faidx ${FASTA_FILE}

# Create a BED file covering the whole genome
cat ${FASTA_FILE}.fai | \
    awk '{print $1"\t0\t"$2}' > whole_genome.bed
```

2. Convert BAM to COV:

```
# Specify the path to the whole-genome BED file in a JSON
echo "{" > annotations_path.json
echo "\"whole_genome\" : \"${PWD}/whole_genome.bed\"" >> annotations_path.json
echo "}" >> annotations_path.json

# Convert BAM to cov.gz using bam2cov
docker run -it --rm -v${WORKING_DIR}:${WORKING_DIR} mobinasri/flagger:v1.1.0 \
    bam2cov --bam ${WORKING_DIR}/${BAM_FILE} \
            --output ${WORKING_DIR}/coverage_file.cov.gz \
            --annotationJson ${WORKING_DIR}/annotations_path.json \
            --threads 16 \
            --baselineAnnotation whole_genome
```

3. Run HMM-Flagger:

```
mkdir -p ${WORKING_DIR}/hmm_flagger_outputs
docker run -it --rm -v${WORKING_DIR}:${WORKING_DIR} mobinasri/flagger:v1.1.0 \
        hmm_flagger \
            --input ${WORKING_DIR}/coverage_file.cov.gz \
            --outputDir ${WORKING_DIR}/hmm_flagger_outputs \
            --alphaTsv /home/programs/config/alpha_optimum_trunc_exp_gaussian_w_4000_n_50.tsv \
            --labelNames Err,Dup,Hap,Col \
            --threads 16
```

Assembly errors were predicted using Flagger, which identifies collapsed, duplicated, and error-prone regions in genome assemblies. Predictions were output in BED format, with regions labeled as DUP (duplication), ERR (error), or COLLAPSE. Only intervals meeting these classification criteria were retained for downstream analysis.

###### 1.12.2 NucFlag-based detection

For NucFlag v0.3.6, PacBio HiFi reads were aligned to the diploid sequence assembly using the following command, retaining only primary alignments:

```
minimap2 -ax lr:hqae --eqx -t {threads} {input_asm} {input_reads} | \
samtools view -F 2308 -u - | \
samtools sort -o {output_bam}
```

NucFlag log files were parsed to extract flagged genomic intervals containing contig coordinates and error-type annotations:

```
nucflag -i {infile} -d {outdir}
```

Regions annotated as heterozygous (HET) or misjoin (MISJOIN) were excluded, as they likely represent biological variation rather than assembly errors.

###### 1.12.3 Reference-based detection

Reference-based evaluation was performed for HG002 assemblies by aligning each assembly to the HG002 Q100 T2T reference and identifying SVs larger than 50 bp. Since both the assemblies and the reference were derived from the same individual, the detected SVs correspond to misassemblies rather than biological variants. The commands used are listed below:

```
minimap2 -cx asm5 -z200000,10000 -t24 --cs hg002v1.1.fasta hap1.asm.fa hap2.asm.fa > asm.paf
sort -k6,6 -k8,8n asm.paf > srt.asm.paf
paftools.js call -f hg002v1.1.fasta srt.asm.paf > srt.asm.vcf
```

###### 1.12.4 Comparison across different approaches

In Supplementary Fig. 5, we analyzed the overlaps among different approaches. Genomic intervals from each pair of methods were compared within their corresponding contig sequences using interval intersection analysis. Two or more intervals were considered overlapping if they shared at least one base pair.

#### 1.13 HG002-specific contig end characterization

In Fig. 5a–c, we characterize contig ends in the HG002 fully phased human genome assemblies to identify regions of assembly difficulty. This analysis integrated alignment-based coordinate mapping with sequence-feature annotation to classify each contig end according to its underlying genomic context.

##### 1.13.1 Alignment and contig end extraction

Each HG002 trio-binning assembly was aligned to its corresponding haplotype-specific reference (HG002 v1.1 maternal or paternal). To improve alignment accuracy, particularly for the sex chromosomes, we specifically aligned the chrX and chrY assemblies to the corresponding chrX and chrY sequences in the reference genome. To further enhance alignment accuracy in highly repetitive regions, we employed a hybrid strategy combining `MashMap` v3.1.3 and `Minimap2` v2.30-r1287. `MashMap` (parameters: `--perc_identity 95 --noSplit`) was first used for coarse chromosome assignment, while `Minimap2` (parameters: `-x asm5 --eqx --cs -N 5`) was subsequently used to generate detailed local alignments. For contigs smaller than 5 kb, for large contigs lacking `MashMap` support, or when `MashMap`-assigned chromosomes were absent from the PAF file, we retained up to the five highest-MAPQ `Minimap2` alignments. From these alignments, 1 kb terminal regions at each contig start and end were extracted to represent potential assembly breakpoints. If the aligned span was shorter than twice the window size (2 × width, 2 kb by default), the entire aligned interval was retained instead of two separate 1 kb windows.

##### 1.13.2 Sequence composition analysis

For each contig end in reference coordinates, nucleotide composition was calculated using `bedtools nuc` v2.31.0, yielding GC content and derived metrics: purine content (GA $= \frac{G+A}{\text{length}}$), pyrimidine content (TC $= \frac{T+C}{\text{length}}$), and AT content. To capture local sequence biases, the maximum value of each metric was computed within a $\pm 10$ kb window centered on each contig end.

##### 1.13.3 Sequence composition analysis

Each contig end was assigned to one of eleven mutually exclusive categories using a priority-based hierarchical classification scheme as follows.

1. Tier 1 – Structural boundaries (highest priority): Contig ends overlapping annotated telomeric regions or located within 200 kb of chromosome ends (minimum distance to start or end) were classified as structural boundaries, encompassing telomeric and subtelomeric sequences.
2. Tier 2 – Satellite sequences:
   - **Alpha satellite:** $\geq$10% overlap with annotated alpha-satellite arrays (CenSat v2.0).
   - **Other satellite:** $\geq$10% overlap with other satellite DNA classes (HSat, $\beta$-satellite).
3. Tier 3 – Segmental duplications (SDs): Contig ends overlapping SD annotations with $\geq$10% reciprocal overlap were included, using SD intervals $\geq$1 kb and $\geq$90% sequence identity. Subclasses were defined as follows:
   - **SD + High GA/TC (80%):** SD overlap combined with extreme purine/pyrimidine strand bias (maximum GA or TC fraction $\geq$0.80 within the contig end region).
   - **High GA/TC (80%):** Extreme strand bias without SD overlap.
   - **SD:** SD overlap without extreme base-composition bias.
4. Tier 4 – Sequence-composition anomalies:
   - **Low complexity:** $\geq$10% overlap with GIAB low-complexity regions (tandem repeats or homopolymers, with $\pm$0 bp slop applied).
   - **High GC ($\geq$65%):** $\geq$10% overlap with GIAB high-GC stratifications; when unavailable, contig ends with calculated maximum GC fraction $\geq$0.75 were included.
   - **High AT ($\geq$70%):** $\geq$10% overlap with GIAB low-GC ($\leq$30%) stratifications; when unavailable, contig ends with calculated maximum AT fraction $\geq$0.80 were included.
5. Tier 5 – Unattributed breaks (lowest priority):

- **Poisson breaks:** Isolated contig ends with $\leq 2$ total ends (including the focal end) within $\pm 100$ kb of the same chromosome, lacking any annotated sequence features. These represent stochastic assembly discontinuities due to coverage or algorithmic fragmentation.
- **Other:** Contig ends not classified by any of the above criteria.

This hierarchical framework ensures that each contig end is attributed to the most specific biological feature available, with "Poisson breaks" serving as a conservative null category after excluding all known difficult-to-assemble sequence contexts. A 10% overlap threshold was used consistently across all annotation layers.

#### 1.14 Assembly gap detection and visualization across all GIAB samples

In Fig. 5d,e, we analyzed ten HiFi and ten ONT diploid genome assemblies from seven GIAB samples, including both fully and partially phased assemblies. For each diploid genome, the two haplotypes were processed independently and annotated with the sample's sex metadata to enable sex-aware analyses.

##### 1.14.1 Reference alignment and chromosome assignment

Contigs were assigned to chromosomes using `MashMap` v3.1.3 against the T2T-CHM13v2.0 reference genome, using a 95% identity threshold to preserve contig integrity. Base-level alignments were then generated with `Minimap2` v2.30 (`-x asm20 --eqx -N 10`) to produce PAF files for downstream filtering and coverage computation.

##### 1.14.2 Sex-aware correction of chrX/Y misassignment

To mitigate chrX/Y cross-mapping, we determined the sex-bearing haplotypes for each sample. For female samples, both haplotypes were treated as X-bearing. For male samples, we computed the number of aligned bases to chrX and chrY for each haplotype using `MashMap`/`Minimap2` outputs, and designated the haplotype with the higher $\mathrm{Y}/(\mathrm{X}+\mathrm{Y})$ ratio as Y-bearing, with the other assigned as X-bearing.

Contigs mapping to the opposite sex chromosome were corrected using the following procedure: (i) checking for existing PAF alignments to the expected chromosome; (ii) if none were present, re-aligning the contig sequence to the CHM13 reference with higher sensitivity (`-x asm20 -N 50`); and (iii) replacing opposite-sex alignments when a valid alignment to the target chromosome was identified. Contigs lacking any detectable alignment to the target chromosome were flagged in `need_patch.list`, but their best-chromosome assignment was still corrected for downstream analyses.

##### 1.14.3 Alignment filtering and multi-layer classification

PAF alignments were filtered to retain only those mapping to the contig's assigned chromosome, keeping primary alignments (`tp:A:P`), requiring MAPQ $\geq$ 20, and restricting to contigs of length $\geq$ 5 kb. We then applied a three-layer contig-quality classification:

- **Layer 1 (high-quality):** overall aligned length $\geq 80\%$ of the contig, and both terminal windows exhibit sufficient coverage (dynamic 1–20 kb windows scaled to contig length).
- **Layer 2 (partial):** overall coverage $\geq 30\%$ but $< 80\%$, or coverage $\geq 80\%$ but with under-covered termini.
- **Layer 3 (low-quality):** overall coverage $< 30\%$; these contigs were excluded from downstream coverage and gap calling.

For Layer 1 contigs, contig ends were localized using an adaptive procedure: beginning at each terminus, a small window (default 3 kb) was shifted inward in increments of up to 500 bp (not exceeding a total of 5 kb) until a minimum coverage threshold was reached. The corresponding reference coordinates were then recorded as the contig start or end positions.

##### 1.14.4 Gap calling

"Covered" reference intervals were defined using Layer 1 and Layer 2 alignments (*coverage strategy: layer1+2*). Overlapping intervals were merged with a 200 bp tolerance. Assembly gaps were defined as the complement of these merged coverage intervals relative to the CHM13 reference (using `bedtools`), and were further size-filtered to retain gaps $\geq$ 1 kb. Gaps located within $\pm 20$ kb of Layer 1 contig ends were flagged as boundary-associated.

###### 1.14.5 Contig-end density

Independently of gap calling, we aggregated reference-mapped contig termini across all samples. For each contig, the strand-aware start and end reference positions on its assigned chromosome were extracted from the filtered PAF alignments. Contig ends from the two haplotypes of the same diploid genome were deduplicated at identical coordinates to avoid double-counting. We then computed position-wise counts of unique samples contributing a contig end, with optional binning (e.g., 20 kb) for visualization.

###### 1.14.6 Sex-aware multi-sample aggregation and normalization

Per-diploid-genome gap sets were aggregated using sex-aware rules:

- **Autosomes:** both haplotypes from all samples.
- **chrX:** males contribute the X-bearing haplotype; females contribute both haplotypes.
- **chrY:** males contribute the Y-bearing haplotype; females are excluded.

Gaps within 1 kb across haplotypes were merged to define consensus regions, and merged intervals shorter than 1 kb were removed. For each consensus gap, we counted the number of overlapping haplotypes and computed a coverage ratio normalized by the expected haplotype denominator for each chromosome:

- autosomes: $2 \times (n_{\text{males}} + n_{\text{females}})$,
- chrX: $(1 \times n_{\text{males}} + 2 \times n_{\text{females}})$,
- chrY: $1 \times n_{\text{males}}$.

where consensus gaps with a ratio $> 0.5$ were designated as "high-confidence recurrent" regions.

###### 1.14.7 Genomic annotations and visualization

We intersected the identified gaps and contig-end density tracks with the following reference annotations: (i) segmental duplications (https://s3-us-west-2.amazonaws.com/human-pangenomics/T2T/CHM13/assemblies/annotation/chm13v2.0_SD.full.bed); (ii) centromeric and satellite regions (https://s3-us-west-2.amazonaws.com/human-pangenomics/T2T/CHM13/assemblies/annotation/chm13v2.0_censat_v2.1.bed); and (iii) their pairwise and combined overlaps. Genome-wide karyotype visualizations integrated per-sample gap heatmaps, contig-end density tracks, and annotation overlays.

###### 1.14.8 Summary statistics

We summarized, for each chromosome and genome-wide, the total number of contig ends, the number of unique contributing samples, the mean and maximum per-position counts, and the distributions of gap coverage ratios, highlighting recurrent assembly hotspots.

### 2 Statistics of ONT standard Simplex and PacBio HiFi data

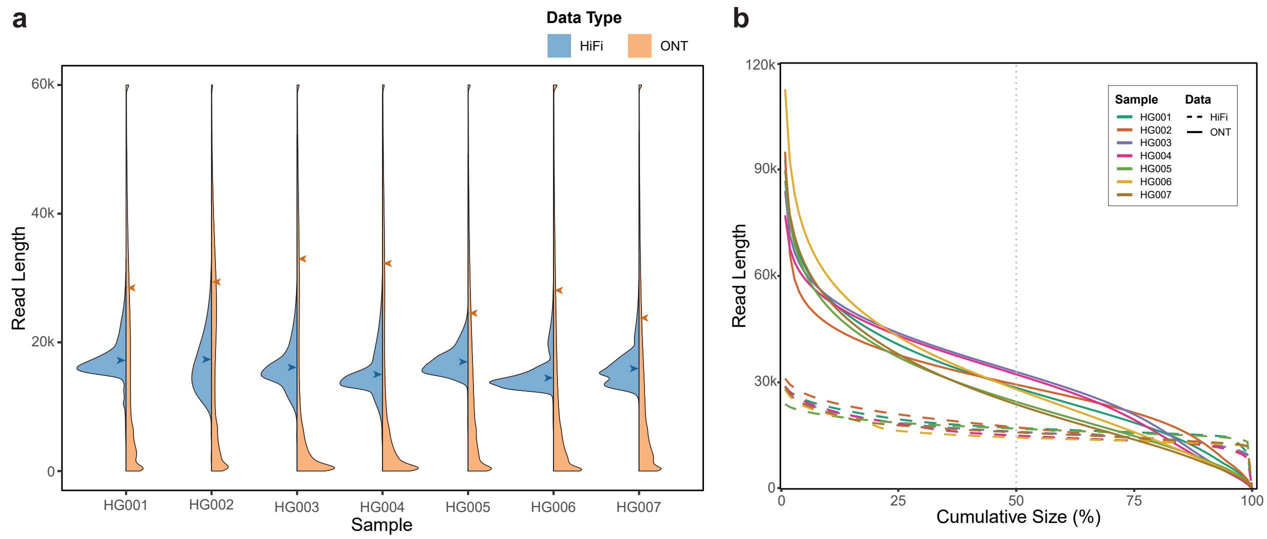


**Supplementary Fig. 1: Comparison of ONT Standard Simplex and PacBio HiFi data for HG001–HG007. (a)** Read length distribution for each sample. Blue and orange arrows indicate the N50 values of ONT Standard Simplex and PacBio HiFi reads, respectively. Read lengths greater than 60 kb are grouped into a single bin. **(b)** Nx plot illustrating read length distributions. For each dataset, the Nx plot shows read lengths sorted from longest to shortest, relative to cumulative read length as a percentage of the total yield.

### 3 Example of ONT assembly errors in long homopolymer regions

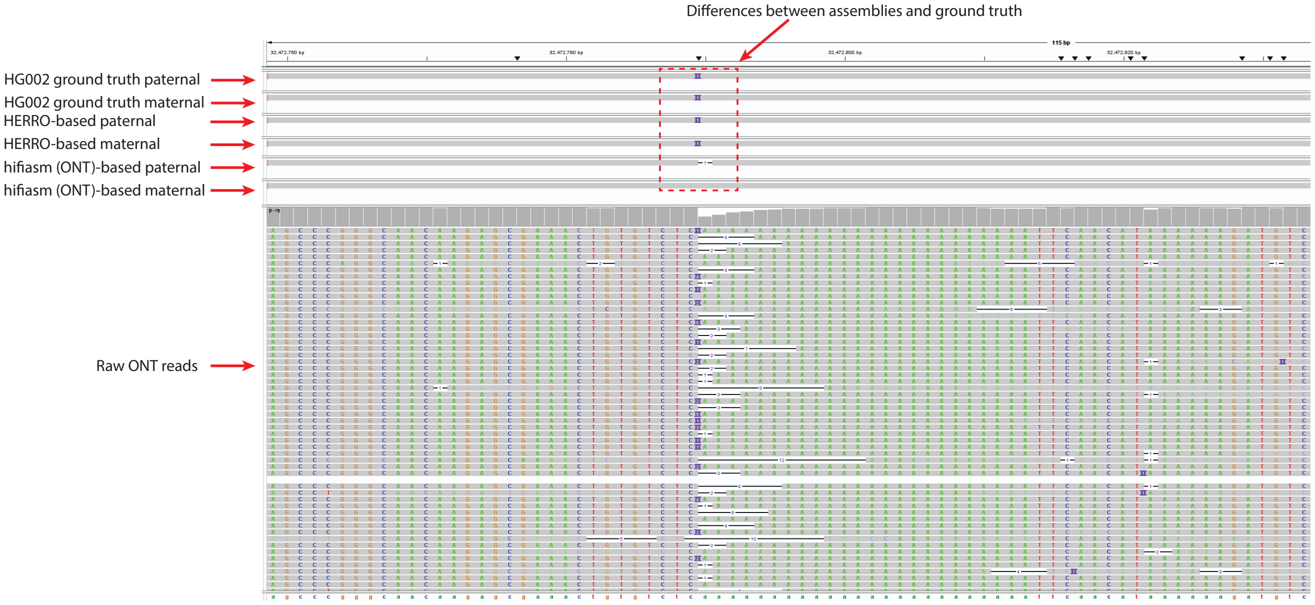


**Supplementary Fig. 2: IGV screenshot showing assembly errors by hifiasm (ONT) within long homopolymer regions.** All ONT assemblies and the HG002 ground truth (HG002 Q100 reference) are aligned to the CHM13 reference genome. Due to sequencing errors in ONT reads around long homopolymer regions (bottom track), hifiasm (ONT) fails to accurately resolve homopolymer lengths.

### 4 Evaluation of phasing errors and QV scores

**Supplementary Table 1:** *k*-mer-based assembly evaluation using yak and Merqury

| Data type | Dataset | Assembler | yak | | | Merqury | | |
|---|---|---|---|---|---|---|---|---|
| | | | QV | switch (%) | hamming (%) | QV | switch (%) | hamming (%) |
| ONT (SUP) standard | HG001 | hifiasm (trio) | 49.08/49.16 | 3.38/1.83 | 4.41/1.21 | 57.21/57.64 | 0.33/0.08 | 0.60/0.10 |
| | | Verkko+HERRO (trio) | 51.38/51.61 | 3.46/1.86 | 4.35/1.27 | 57.43/57.64 | 0.29/0.03 | 0.48/0.10 |
| | | hifiasm (dual) | 49.12/49.13 | / | / | 57.24/57.69 | / | / |
| | HG002 | hifiasm (trio) | 47.54/47.12 | 1.59/2.52 | 1.22/1.69 | 51.50/51.50 | 0.08/0.11 | 0.11/0.16 |
| | | Verkko+HERRO (trio) | 48.97/48.48 | 1.62/2.61 | 1.23/1.73 | 51.67/51.57 | 0.06/0.07 | 0.08/0.10 |
| | | hifiasm (dual) | 47.33/47.29 | / | / | 51.49/51.64 | / | / |
| | HG003 | hifiasm (dual) | 48.20/47.69 | / | / | 64.69/64.87 | / | / |
| | HG004 | hifiasm (dual) | 48.53/48.61 | / | / | 64.46/63.13 | / | / |
| | HG005 | hifiasm (trio) | 48.28/48.42 | 0.88/2.08 | 0.57/1.46 | 49.78/49.89 | 0.05/0.09 | 0.06/0.16 |
| | | Verkko+HERRO (trio) | 50.64/50.68 | 0.91/2.10 | 0.56/1.43 | 49.87/49.93 | 0.01/0.03 | 0.02/0.04 |
| | | hifiasm (dual) | 48.45/48.35 | / | / | 49.80/49.89 | / | / |
| | HG006 | hifiasm (dual) | 47.88/47.64 | / | / | 63.42/61.87 | / | / |
| | HG007 | hifiasm (dual) | 47.49/47.83 | / | / | 59.57/61.45 | / | / |
| ONT (SUP) ultra-long | HG002 | hifiasm (trio) | 46.12/45.66 | 1.69/2.71 | 1.26/1.75 | 50.76/50.44 | 0.06/0.06 | 0.07/0.09 |
| | | Verkko+HERRO (trio) | 47.79/47.37 | 1.68/2.44 | 1.25/1.59 | 51.08/51.03 | 0.07/0.06 | 0.07/0.09 |
| | HG02818 | hifiasm (trio) | 45.57/45.59 | 3.18/3.16 | 1.94/2.20 | 54.42/54.17 | 0.04/0.05 | 0.04/0.25 |
| | | Verkko+HERRO (trio) | 47.46/47.48 | 3.21/3.12 | 1.98/2.04 | 55.10/55.14 | 0.05/0.06 | 0.06/0.06 |
| | Arabidopsis | hifiasm | 43.93 | / | / | 41.73 | / | / |
| | | Verkko+HERRO | 45.30 | / | / | 33.35 | / | / |
| | Tomato | hifiasm | 42.48 | / | / | 45.54 | / | / |
| | | Verkko+HERRO | 43.18 | / | / | 45.40 | / | / |
| | Zebrafish | hifiasm | 43.06 | / | / | 56.95 | / | / |
| | | Verkko+HERRO | 43.59 | / | / | 51.05 | / | / |
| HiFi | HG001 | hifiasm (trio) | 50.21/49.97 | 2.65/1.14 | 4.04/0.86 | 51.86/51.75 | 0.07/0.08 | 0.14/0.11 |
| | | hifiasm (dual) | 50.12/50.02 | / | / | 51.90/51.78 | / | / |
| | HG002 | hifiasm (trio) | 53.78/53.00 | 0.80/0.96 | 0.77/0.70 | 51.86/51.75 | 0.07/0.08 | 0.14/0.11 |
| | | hifiasm (dual) | 53.03/53.81 | / | / | 51.67/51.88 | / | / |
| | HG003 | hifiasm (dual) | 51.75/52.39 | / | / | 56.61/62.83 | / | / |
| | HG004 | hifiasm (dual) | 53.22/54.29 | / | / | 60.30/62.12 | / | / |
| | HG005 | hifiasm (trio) | 53.59/53.39 | 0.29/0.73 | 0.21/0.62 | 49.71/49.74 | 0.03/0.06 | 0.04/0.10 |
| | | hifiasm (dual) | 53.48/53.52 | / | / | 49.63/49.81 | / | / |
| | HG006 | hifiasm (dual) | 55.56/55.97 | / | / | 54.53/63.54 | / | / |
| | HG007 | hifiasm (dual) | 53.42/54.55 | / | / | 57.44/58.45 | / | / |
| ONT (HAC) standard | HG001 | hifiasm (trio) | 45.68/45.66 | 3.94/2.39 | 4.86/1.62 | 53.19/53.10 | 0.41/0.15 | 0.74/0.23 |
| | | hifiasm (dual) | 45.62/45.71 | / | / | 52.98/53.18 | / | / |
| | HG002 | hifiasm (trio) | 44.43/44.01 | 1.98/3.36 | 1.50/2.25 | 49.21/49.06 | 0.12/0.18 | 0.16/0.25 |
| | | hifiasm (dual) | 44.07/44.35 | / | / | 49.11/49.13 | / | / |
| | HG003 | hifiasm (dual) | 45.28/44.92 | / | / | 56.86/57.68 | / | / |
| | HG004 | hifiasm (dual) | 45.73/45.78 | / | / | 57.68/57.68 | / | / |
| | HG005 | hifiasm (trio) | 45.06/45.15 | 1.22/2.82 | 0.83/1.98 | 48.67/48.68 | 0.09/0.17 | 0.15/0.24 |
| | | hifiasm (dual) | 45.13/45.10 | / | / | 48.71/48.63 | / | / |
| | HG006 | hifiasm (dual) | 44.27/44.13 | / | / | 55.13/55.48 | / | / |
| | HG007 | hifiasm (dual) | 44.38/44.66 | / | / | 53.69/55.35 | / | / |

The phasing switch error rate refers to the proportion of adjacent haplotype-specific marker pairs originating from different haplotypes, while the phasing hamming error rate represents the percentage of haplotype-specific markers that are incorrectly phased. Only fully phased trio-binning assemblies were evaluated for phasing accuracy. All assemblies were assessed for QV. For *Linum usitatissimum* (flax), we did not perform *k*-mer-based assembly evaluation because short-read data from the same individual are not available.

### 5 Assembly results using ONT Simplex reads with the HAC basecalling model

**Supplementary Table 2:** Statistics of hifiasm assemblies using ONT standard Simplex reads (HAC basecalling model)

| Phased | Sample | Wall time (h) | T2T count | | Multicopy genes retained (%) | N50 (Mb) | | QV |
|---|---|---|---|---|---|---|---|---|
| | | | Scaffold | Contig | | Scaffold | Contig | |
| full (trio) | HG001 | 13.7 | 12/10 | 8/5 | 92.4/92.6 | 134.5/127.4 | 102.1/109.3 | 45.7/45.7 |
| | HG002 | 7.5 | 13/13 | 7/7 | 91.5/95.0 | 135.0/136.5 | 132.2/103.3 | 44.4/44.0 |
| | HG005 | 9.9 | 10/11 | 4/2 | 92.7/93.4 | 107.0/107.7 | 103.2/90.2 | 45.1/45.1 |
| partial (dual) | HG001 | 13.5 | 13/15 | 8/9 | 94.5/90.5 | 135.4/135.7 | 111.5/105.0 | 45.6/45.7 |
| | HG002 | 7.4 | 14/11 | 7/5 | 93.1/93.4 | 135.7/135.0 | 101.3/101.3 | 44.1/44.4 |
| | HG003 | 17.0 | 13/12 | 9/7 | 94.9/96.1 | 134.9/134.0 | 130.5/99.6 | 45.3/44.9 |
| | HG004 | 13.5 | 10/10 | 7/3 | 92.8/94.2 | 111.0/135.6 | 100.4/90.9 | 45.7/45.8 |
| | HG005 | 9.8 | 0/4 | 0/4 | 92.4/93.4 | 100.9/104.7 | 83.6/88.5 | 45.1/45.1 |
| | HG006 | 9.9 | 14/12 | 9/5 | 94.1/95.0 | 134.4/109.0 | 96.5/87.3 | 44.3/44.1 |
| | HG007 | 13.3 | 11/11 | 5/6 | 89.6/96.4 | 111.5/146.4 | 98.1/134.6 | 44.4/44.7 |

### 6 Comparison of hifiasm (ONT) and Napu (Shasta) assemblers

We compared hifiasm (ONT) with an existing ONT-based haplotype-resolved assembler, Napu (Nanopore Analysis Pipeline) (https://github.com/nanoporegenomics/napu_wf). Napu is built upon Shasta and supports both genome assembly and assembly-based variant calling only with ONT reads. We generated hifiasm (ONT) assemblies for three human samples (HG002, HG00733, and HG02723) and compared them to Napu assemblies using the same ONT standard Simplex read dataset for each sample. Napu assemblies were produced by the Napu developers (https://s3-us-west-2.amazonaws.com/human-pangenomics/index.html?prefix=publications/Napu_paper_ONT_Coriell_SingleFC_2023/).

As shown in Supplementary Table 3, the contig N50 values of the hifiasm (ONT) assemblies are lower than those shown in Table 1 due to lower read coverage. Compared to Napu, hifiasm (ONT) achieves substantially higher contiguity (contig N50) and better resolution of repetitive regions, as indicated by the retained multicopy genes. The smaller size of Napu assemblies also indicates that it collapses much more repetitive regions.

**Supplementary Table 3:** Assembly statistics of hifiasm and Napu (Shasta) using ONT standard Simplex reads

| Sample | Approach | Contig N50 (Mb) | | Multicopy genes retained (%) | | Size (Gb) | |
|---|---|---|---|---|---|---|---|
| | | haplotype 1 | haplotype 2 | haplotype 1 | haplotype 2 | haplotype 1 | haplotype 2 |
| HG002 (42x) | hifiasm | 80.69 | 74.45 | 94.10 | 92.49 | 3.05 | 2.95 |
| | Napu (Shasta) | 14.92 | 14.91 | 64.62 | 64.67 | 2.93 | 2.93 |
| HG00733 (43x) | hifiasm | 82.85 | 80.67 | 94.01 | 95.34 | 3.03 | 3.00 |
| | Napu (Shasta) | 25.33 | 25.34 | 63.24 | 61.91 | 2.90 | 2.90 |
| HG02723 (38x) | hifiasm | 57.38 | 59.78 | 93.06 | 91.01 | 3.03 | 3.02 |
| | Napu (Shasta) | 19.10 | 19.08 | 61.77 | 62.01 | 2.88 | 2.88 |

### 7 Assembly results of the flax genome

**Supplementary Table 4:** Statistics of flax genome assemblies using ONT standard Simplex reads (SUP basecalling model)

| Data type | Assembler | Wall time (h) | T2T count | | N50 (Mb) | |
|---|---|---|---|---|---|---|
| | | | Scaffold | Contig (%) | Contig | Scaffold |
| ONT standard | hifiasm | 9.2 | 12 | 12 | 32.14 | 32.14 |
| (120x) | Verkko+HERRO | 26.27 | 2 | 2 | 13.99 | 13.99 |

Flax (*Linum usitatissimum* L.) is a nearly homozygous species with 15 chromosomes. Both hifiasm (ONT) and Verkko were run using their haploid assembly modules. The scaffolding approaches implemented in these assemblers are not applicable to haploid genome assembly.

### 8 Annotated immunological genes per human genome assembly

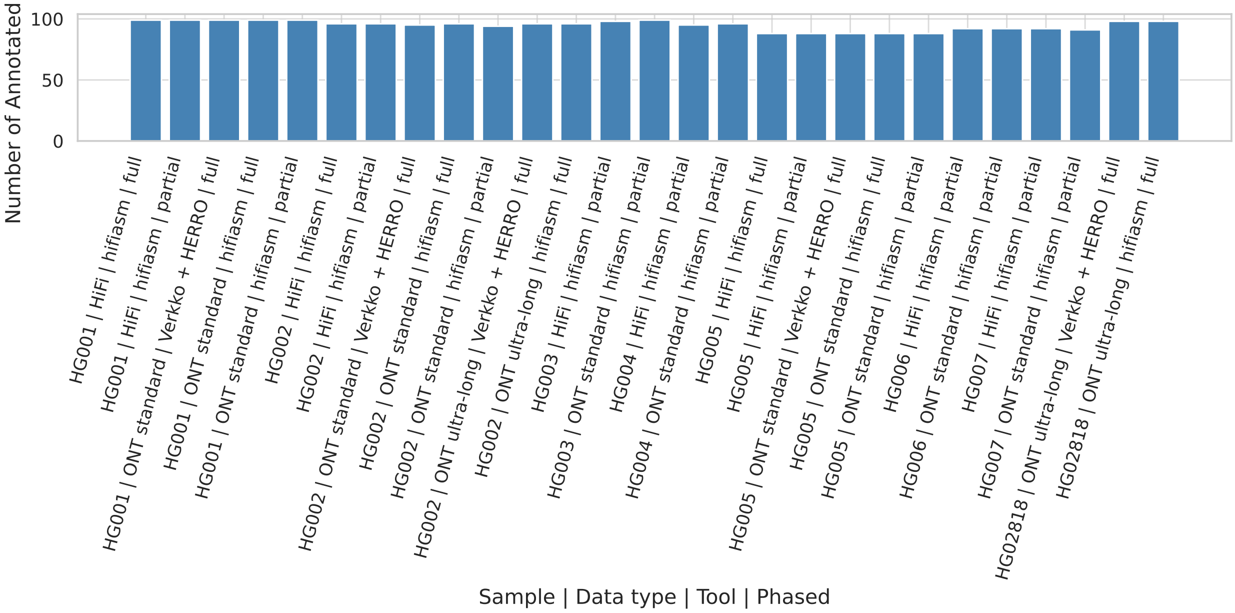


**Supplementary Fig. 3: Comparison of the number of annotated HLA and KIR genes across different human genome assemblies.** Gene annotation was performed using Immuannot, which aligns reference gene sequences to each assembly. For a given assembly, a gene may be unreported if its sequence diverges substantially from the reference.

### 9 Evaluation of assembly errors

**Supplementary Table 5:** Structural variant statistics from HG002 phased assemblies compared to the HG002 Q100 reference genome

| Data | Approach | Contig N50 (Mb) | SV total length (Mb) | # SV (no merge) | # SV (merge by distance) | | |
|---|---|---|---|---|---|---|---|
| | | | | | 1,000 bp | 5,000 bp | 10,000 bp |
| ONT ultra-long | hifiasm | 146.34 / 143.79 | 0.4 | 184 | 160 | 146 | 135 |
| | Verkko + HERRO | 97.49 / 111.04 | 1.5 | 751 | 679 | 399 | 269 |
| ONT standard | hifiasm | 131.52 / 143.80 | 1.9 | 402 | 353 | 246 | 191 |
| | Verkko + HERRO | 52.29 / 45.99 | 3.7 | 934 | 837 | 727 | 661 |
| PacBio HiFi | hifiasm | 78.70 / 63.10 | 3.6 | 772 | 657 | 524 | 450 |

Each assembly consists of two sets of sequences representing the paternal and maternal haplotypes. The two numbers in each cell correspond to the metrics for the two haplotypes. These SVs are often misassemblies, as both the assembly and reference originate from the same individual.

### 10 Distribution of misassemblies across HG002 assemblies

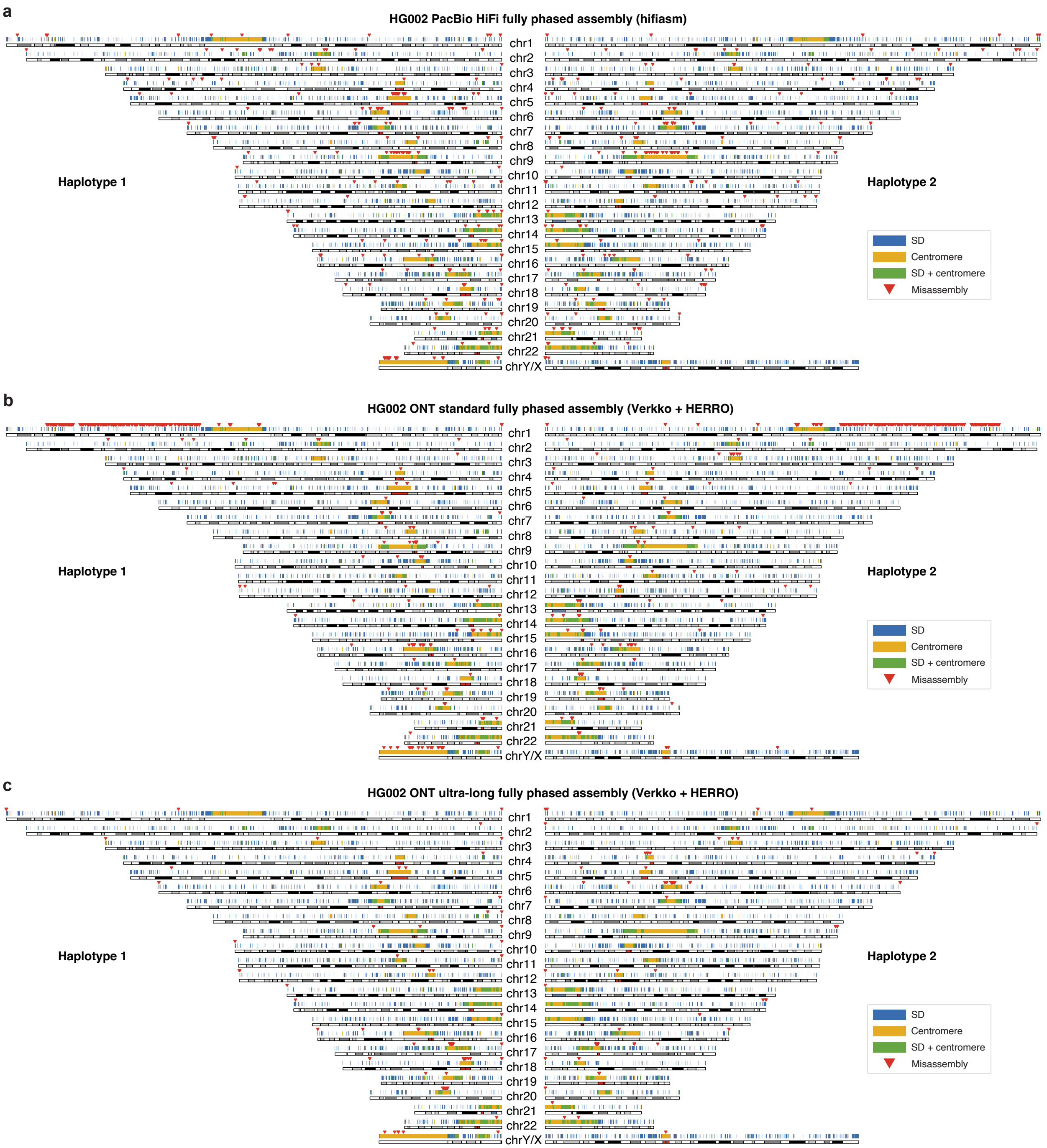


**Supplementary Fig. 4: Genome-wide distribution of misassemblies in HG002 assemblies.** Only misassemblies larger than 50 bp were counted, and all misassemblies were accurately assessed against the HG002 Q100 reference genome. Segmental duplication and centromere annotations are shown in blue and yellow, with overlapping regions highlighted in green and misassembly sites marked by red triangles. **(a)** Misassemblies in the HG002 PacBio HiFi fully phased assembly generated by hifiasm. **(b)** Misassemblies in the HG002 ONT standard fully phased assembly produced by Verkko + HERRO. **(c)** Misassemblies in the HG002 ONT ultra-long fully phased assembly produced by Verkko + HERRO.

### 11 Comparison of misassembly detection across different approaches

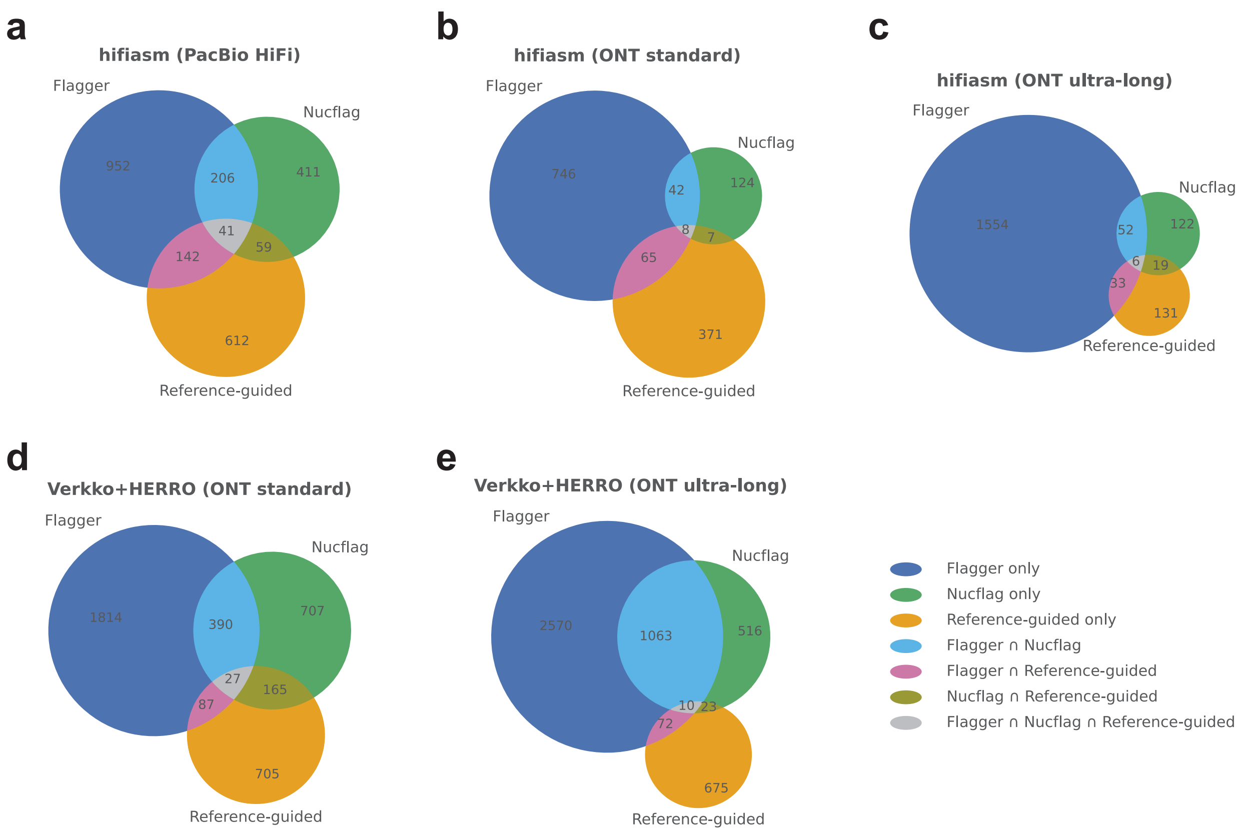


**Supplementary Fig. 5: Number of overlapping misassemblies identified by Flagger, Nucflag, and reference-guided analyses in HG002 assemblies.** Only misassemblies larger than 50 bp were included. Reference-guided misassemblies were evaluated against the HG002 Q100 ground truth, while Flagger and Nucflag represent reference-free methods that align reads back to assemblies. **(a)** HG002 PacBio HiFi fully phased assembly generated by hifiasm. **(b)** HG002 ONT standard fully phased assembly produced by hifiasm (ONT). **(c)** HG002 ONT ultra-long fully phased assembly produced by hifiasm (ONT). **(d)** HG002 ONT standard fully phased assembly produced by Verkko + HERRO. **(e)** HG002 ONT ultra-long fully phased assembly produced by Verkko + HERRO.

### 12 Assembly-based variant calling results for HG002

**Supplementary Table 6:** HG002 fully phased assembly variant calling compared with the HG002 benchmark (SUP basecalling model)

| Data type | Assembler | Region | SNP (%) | | | INDEL (%) | | | ALL (%) | | |
|---|---|---|---|---|---|---|---|---|---|---|---|
| | | | Precision | Recall | F1 | Precision | Recall | F1 | Precision | Recall | F1 |
| ONT standard | hifiasm | homopolymer | 96.56 | 96.00 | 96.28 | 47.80 | 64.55 | 54.93 | 74.47 | 84.46 | 79.15 |
| | | non-homopolymer | 97.72 | 97.04 | 97.38 | 94.77 | 93.57 | 94.17 | 97.39 | 96.69 | 97.04 |
| | Verkko + HERRO | homopolymer | 96.83 | 96.23 | 96.53 | 55.71 | 65.54 | 60.23 | 79.68 | 84.97 | 82.24 |
| | | non-homopolymer | 97.75 | 97.12 | 97.43 | 95.65 | 94.68 | 95.16 | 97.51 | 96.87 | 97.19 |
| ONT ultra-long | hifiasm | homopolymer | 96.22 | 95.80 | 96.01 | 41.67 | 61.31 | 49.61 | 70.41 | 83.15 | 76.25 |
| | | non-homopolymer | 97.72 | 97.08 | 97.40 | 93.44 | 93.35 | 93.39 | 97.25 | 96.70 | 96.98 |
| | Verkko + HERRO | homopolymer | 96.59 | 96.06 | 96.32 | 50.96 | 63.24 | 56.44 | 77.01 | 84.02 | 80.36 |
| | | non-homopolymer | 97.70 | 97.13 | 97.41 | 93.80 | 93.44 | 93.62 | 97.27 | 96.76 | 97.02 |

Assemblies were aligned to GRCh38, and variants were called using dipcall. Variant calling results was evaluated using hap.py.

### 13 Gene-level resolution in HG002 assemblies

**Supplementary Table 7:** Summary of gene resolution in HG002 fully phased assemblies relative to the HG002 Q100 reference genome

| Data | Approach | Contig N50 (Mb) | # Single-copy resolved | # Multicopy resolved | # False duplicated | # Partially resolved | # Unresolved | | |
|---|---|---|---|---|---|---|---|---|---|
| | | | | | | | $> 50\%$ | 10–50% | $\leq 10\%$ |
| HG002 Ref | mannually curated | 146.8 / 154.3 | 34114/35754 | 1203/1276 | NA | NA | NA | NA | NA |
| PacBio HiFi | hifiasm | 78.7 / 63.1 | 33876/35403 | 1176/1265 | 32/23 | 7/5 | 10/26 | 5/8 | 184/289 |
| ONT (SUP) | hifiasm | 131.5 / 143.8 | 34092/35731 | 1199/1272 | 8/7 | 0/0 | 2/1 | 0/0 | 12/15 |
| standard | Verkko + HERRO | 52.3 / 46.0 | 34001/35631 | 1188/1260 | 70/74 | 3/16 | 6/4 | 3/1 | 31/28 |
| ONT (SUP) | hifiasm | 146.3 / 143.8 | 34078/35698 | 1198/1274 | 13/34 | 0/0 | 3/2 | 1/0 | 19/20 |
| ultra-long | Verkko + HERRO | 97.5 / 111.0 | 34090/35733 | 1197/1274 | 11/9 | 0/0 | 0/2 | 0/0 | 13/10 |

Each assembly consists of two sets of sequences representing the paternal and maternal haplotypes. The two numbers in each cell correspond to the metrics for the two haplotypes, respectively. For each assembly or genome, genes were identified by aligning cDNA sequences with a sequence identity of at least 99%. 'HG002 Ref' refers to the T2T Q100 reference genome for HG002, whose aligned genes were used as the ground truth. Gene completeness was evaluated by comparing the genes identified in each assembly with those in the HG002 reference. 'Single-copy resolved' denotes the number of single-copy genes in the HG002 reference that remain single-copy in the assembly. 'Multicopy resolved' denotes the number of multi-copy genes in the HG002 reference that remain multi-copy in the assembly. 'False duplications' represent single-copy genes in the HG002 reference that appear as multi-copy in the assembly. 'Partially resolved' refers to genes that are fully present but fragmented across multiple pieces in the assembly. 'Unresolved' ($> 50\%$, 10–50%, $\leq 10\%$) indicates genes that cannot be completely identified in the assembly, where the aligned length covers more than 50%, between 10% and 50%, or less than 10% of the full gene length, respectively.

### 14 Sequencing datasets used in this study

**Supplementary Table 8:** Sequencing datasets used in this study (full links)

| Sample | Data type | Download link/accession |
|---|---|---|
| HG001 | ONT standard (SUP) | s3://ont-open-data/giab_2025.01/basecalling/sup/HG001 |
| HG001 | ONT standard (HAC) | s3://ont-open-data/giab_2025.01/basecalling/hac/HG001 |
| HG001 | PacBio HiFi | https://ftp.ncbi.nlm.nih.gov/ReferenceSamples/giab/data/NA12878/HudsonAlpha_PacBio_CCS/ |
| HG001 | Illumina short reads | ERR194147 (HG001), ERR194160 (paternal), ERR194161 (maternal) |
| HG002 | ONT standard (SUP) | s3://ont-open-data/giab_2025.01/basecalling/sup/HG002/PAW70337 |
| HG002 | ONT standard (HAC) | s3://ont-open-data/giab_2025.01/basecalling/hac/HG002/PAW70337 |
| HG002 | ONT ultra-long | https://s3-us-west-2.amazonaws.com/human-pangenomics/index.html?prefix=submissions/5b73fa0e-658a-4248-b2b8-cd16155bc157--UCSC_GIAB_R1041_nanopore/HG002_R1041_UL/dorado/v0.4.0_wMods/*ULCIR*.bam https://s3-us-west-2.amazonaws.com/human-pangenomics/index.html?prefix=submissions/5b73fa0e-658a-4248-b2b8-cd16155bc157--UCSC_GIAB_R1041_nanopore/HG002_R1041_UL/dorado/v0.4.0_wMods/*ULNEB*.bam |
| HG002 | PacBio HiFi | https://ftp.ncbi.nlm.nih.gov/ReferenceSamples/giab/data/AshkenazimTrio/HG002_NA24385_son/PacBio_HiFi-Revio_20231031/ |
| HG002 | Illumina short reads | https://s3-us-west-2.amazonaws.com/human-pangenomics/index.html?prefix=working/HPRC_PLUS/HG002/raw_data/Illumina/ |
| HG003 | ONT standard (SUP/HAC) | s3://ont-open-data/giab_2025.01/basecalling/{sup,hac}/HG003 |
| HG003 | PacBio HiFi | https://ftp.ncbi.nlm.nih.gov/ReferenceSamples/giab/data/AshkenazimTrio/HG003_NA24149_father/PacBio_HiFi-Revio_20231031/ |

(continued on next page)

*(continued from previous page)*

| Sample | Data Type | Download link/accession |
|---|---|---|
| | | https://ftp.ncbi.nlm.nih.gov/ReferenceSamples/giab/data/AshkenazimTrio/HG003_NA24149_father/PacBio_CCS_Google_15kb/ |
| HG003 | Illumina short reads | https://s3-us-west-2.amazonaws.com/human-pangenomics/index.html?prefix=working/HPRC_PLUS/HG002/raw_data/Illumina/parents/HG003/ |
| HG004 | ONT standard (SUP/HAC) | s3://ont-open-data/giab_2025.01/basecalling/{sup,hac}/HG004 |
| HG004 | PacBio HiFi | https://ftp.ncbi.nlm.nih.gov/ReferenceSamples/giab/data/AshkenazimTrio/HG004_NA24143_mother/PacBio_HiFi-Revio_20231031/ <br> https://ftp.ncbi.nlm.nih.gov/ReferenceSamples/giab/data/AshkenazimTrio/HG004_NA24143_mother/PacBio_CCS_Google_15kb/ |
| HG004 | Illumina short reads | https://s3-us-west-2.amazonaws.com/human-pangenomics/index.html?prefix=working/HPRC_PLUS/HG002/raw_data/Illumina/parents/HG004/ |
| HG005 | ONT standard (SUP/HAC) | s3://ont-open-data/giab_2025.01/basecalling/{sup,hac}/HG005 |
| | PacBio HiFi | https://ftp.ncbi.nlm.nih.gov/ReferenceSamples/giab/data/ChineseTrio/HG005_NA24631_son/PacBio_CCS_15kb_20kb_chemistry2/uBAMs/ |
| | Illumina short reads | https://s3-us-west-2.amazonaws.com/human-pangenomics/index.html?prefix=working/HPRC_PLUS/HG005/raw_data/Illumina/ |
| HG006 | ONT standard (SUP/HAC) | s3://ont-open-data/giab_2025.01/basecalling/{sup,hac}/HG006 |
| | PacBio HiFi | https://ftp.ncbi.nlm.nih.gov/ReferenceSamples/giab/data/ChineseTrio/HG006_NA24694-huCA017E_father/PacBio_CCS_15kb_20kb_chemistry2/uBAMs/ <br> https://ftp.ncbi.nlm.nih.gov/ReferenceSamples/giab/data/ChineseTrio/HG006_NA24694-huCA017E_father/PacBio_HiFi_Google/ |
| | Illumina short reads | https://s3-us-west-2.amazonaws.com/human-pangenomics/index.html?prefix=working/HPRC_PLUS/HG005/raw_data/Illumina/parents/HG006/ |
| HG007 | ONT standard (SUP/HAC) | s3://ont-open-data/giab_2025.01/basecalling/{sup,hac}/HG007 |
| | PacBio HiFi | https://ftp.ncbi.nlm.nih.gov/ReferenceSamples/giab/data/ChineseTrio/HG007_NA24695-hu38168_mother/PacBio_CCS_15kb_20kb_chemistry2/uBAMs/ <br> https://ftp.ncbi.nlm.nih.gov/ReferenceSamples/giab/data/ChineseTrio/HG007_NA24695-hu38168_mother/PacBio_HiFi_Google/ |
| | Illumina short reads | https://s3-us-west-2.amazonaws.com/human-pangenomics/index.html?prefix=working/HPRC_PLUS/HG005/raw_data/Illumina/parents/HG007/ |
| HG02818 | ONT ultra-long | https://s3-us-west-2.amazonaws.com/human-pangenomics/index.html?prefix=working/HPRC_PLUS/HG02818/raw_data/nanopore/dorado0.7.2_sup4.1.0_5mCG_5hmCG/ |
| HG02818 | Illumina short reads | https://s3-us-west-2.amazonaws.com/human-pangenomics/index.html?prefix=working/HPRC_PLUS/HG02818/raw_data/Illumina/ |
| *Arabidopsis thaliana* | ONT ultra-long | SRR29061597 |
| *Arabidopsis thaliana* | Illumina short reads | https://ngdc.cncb.ac.cn/gsa/browse/CRA005350 |
| *Danio rerio* | ONT ultra-long | https://genomeark.s3.amazonaws.com/index.html?prefix=species/Danio_rerio/fDanRer17/genomic_data/ont/pod5/ <br> https://genomeark.s3.amazonaws.com/index.html?prefix=species/Danio_rerio/fDanRer17/genomic_data/ont/fast5/ |
| *Danio rerio* | Illumina short reads | https://www.ncbi.nlm.nih.gov/sra?linkname=bioproject_sra_all&from_uid=1029986 |

(continued on next page)

*(continued from previous page)*

| Sample | Data Type | Download link/accession |
| --- | --- | --- |
| *Solanum lycopersicum* | ONT ultra-long | https://obj.umiacs.umd.edu/marbl_publications/duplex/Solanum_lycopersicum_heinz1706/UL/R10.4_40x.noduplex.fastq.gz |
| *Solanum lycopersicum* | Illumina short reads | https://ngdc.cncb.ac.cn/gsa/browse/CRA003995/CRX232533 |
| *Linum usitatissimum* | ONT standard | SRR31124331 |
| HG002, HG00733, HG02723 (Napu) | ONT R10 reads | https://s3-us-west-2.amazonaws.com/human-pangenomics/index.html?prefix=publications/Napu_paper_ONT_Coriell_SingleFC_2023/HG0*_R10/reads/*.bam |
